# Microbial carbon metabolism is linked to organic matter chemistry across soil systems

**DOI:** 10.64898/2026.01.20.700515

**Authors:** Daniel Wasner, Oliver Lechtenfeld, Jan Kaesler, Sebastian Doetterl, Meret Aeppli

**Affiliations:** Soil Biogeochemistry Laboratory, Swiss Federal Institute of Technology Lausanne (EPFL), Sion, Switzerland; Department of Environmental Analytical Chemistry, Research Group BioGeoOmics, Helmholtz Centre for Environmental Research–UFZ, Leipzig, Germany; Soil Resources, Department of Environmental System Science, ETH Zurich, Zurich, Switzerland

**Keywords:** microbial growth, microbial respiration, microbial metabolism, carbon use efficiency, soil carbon turnover, organic matter composition, organic matter chemistry, carbon oxidation state

## Abstract

Soil microbial growth and respiration play a critical role for soil organic carbon dynamics. Yet, we lack understanding of the main controls of soil microbial carbon metabolism at large scales. Here, we investigated whether and how the chemical composition of microbially available organic matter affects soil microbial carbon metabolism across soil systems. We linked soil microbial growth and respiration rates as well as carbon use efficiency (quantified with ^18^O stable isotope probing) to the chemical composition of extractable organic matter (characterized with reversed-phase liquid chromatography coupled to Fourier-transform ion cyclotron resonance mass spectrometry) along a geoclimatic gradient of 33 Chilean temperate grassland soils. We found that biomass-normalized rates of growth and respiration were primarily positively linked to aliphatics such as carbohydrate-, proteinaceous- and amino sugar-like compounds, and secondarily to unsaturated lignin-like compounds. Respiration was positively linked to compounds with carbon in a reduced oxidation state, suggesting carbon-conserving catabolism, while growth appeared unrelated to the oxidation state of carbon. This suggests that other mechanisms than mere energetic constraints control microbial growth rates in aerated soils. Our findings demonstrate that information on the chemical composition of bioavailable organic matter can provide insights into the processes that govern the fate of carbon across different ecosystems.

**Key points:**

- We investigated if bulk soil microbial growth (^18^O stable isotope probing) and respiration is linked to the chemical composition of extractable organic matter (LC-FT-ICR MS) along a geoclimatic gradient of temperate grasslands.
- Higher rates of microbial carbon turnover were positively linked to aliphatic and unsaturated compounds.
- Specific (i.e., biomass-normalized) respiration was positively linked to compounds with carbon in a reduced oxidation state, suggesting carbon-conserving catabolism.
- Specific growth was unrelated to the oxidation states of substrate carbon, suggesting that soil microbial substrate use for anabolism may not be determined by direct energetic constraints.

## 1. Introduction

Soil microbial carbon turnover is an important turnstile in the terrestrial carbon (C) cycle. Terrestrial ecosystems have a net uptake of 2 – 4 Gt C per year from the atmosphere through photosynthetic conversion of CO_2_ into organic matter (Friedlingstein et al., 2023). Ultimately, much of this plant organic matter enters the soil, either as exudates or dead plant biomass. In soil, microbes transform the majority of this organic matter through their metabolism. In aerated soils, heterotrophic respiration is the dominant catabolic process by which microbes extract energy from organic substrates. On the one hand, heterotrophic respiration leads to the loss of soil organic carbon as CO_2_, which is the main source of C loss from soils (Kopittke et al., 2024). On the other hand, microbial carbon transformation through anabolic processes such as growth - with the subsequent formation of necromass - are a major pathway for the formation of stable mineral-associated organic matter (Liang, Schimel and Jastrow, 2017; Kästner et al., 2021; Sokol et al., 2022), which can become stabilized and stay in soil for years to millenia.

However, despite the knowledge that microbial growth and respiration are central to soil organic carbon turnover, numerical models still struggle to match empirical observations of soil respiration and soil organic carbon stocks at regional to global scales (Sulman et al., 2018; Georgiou et al., 2021; Hashimoto, Ito and Nishina, 2023). One source of uncertainty is the representation of soil microbial carbon metabolism at the macroscale (Wieder et al., 2015; Chandel, Jiang and Luo, 2023) - which mechanisms and controls of microbial carbon metabolism need to be scaled from micro- to macroscale? In a previous study, we found that potential (i.e., temperature- and moisture-controlled) bulk microbial carbon metabolism along a large geoclimatic grassland gradient was strongly affected by microbial community composition (Wasner et al., 2024b). However, parts of the variation in specific respiration (27%), specific growth (53%) and CUE (35%) still remained unexplained in that study.

Thermodynamic aspects of soil microbial metabolism have recently received increased attention as potential controls for microbial carbon turnover because the majority of soil microbes are chemoorganoheterotrophs, which means that they derive both energy as well as carbon from organic matter (Bajracharya et al., 2022; Gunina and Kuzyakov, 2022; Kästner et al., 2024; Chakrawal et al., 2025). Because soil organic matter is a complex mixture of compounds with diverse degrees of oxidation, functional groups and elemental stoichiometry (Simpson and Simpson, 2012; Lehmann and Kleber, 2015), the chemical composition of microbially available soil organic matter could therefore affect microbial carbon turnover (Boye et al., 2018; Jones et al., 2018; Chakrawal et al., 2022; Gunina and Kuzyakov, 2022). In well-aerated soils, the nominal oxidation state of carbon (NOSC) of a compound can serve as a simplified proxy for the amount of energy that can be obtained per mole carbon via complete oxidation of an organic compound with O_2_ to CO_2_ (LaRowe and Van Cappellen, 2011; Kästner et al., 2024). Oxidation of a more oxidised compound (i.e, with higher NOSC) releases more energy per electron, but fewer electrons are available per carbon (LaRowe and Van Cappellen, 2011). In contrast, the oxidation of a compound with a lower NOSC releases more electrons per carbon, and is therefore more carbon-conserving (Gunina and Kuzyakov, 2022) allowing a higher CUE - at the expense of energy (Chakrawal et al., 2022). Energy-limited microbes may metabolize at a lower CUE due to oxidation of more oxidised compounds, while carbon-limited microbes may increase CUE through oxidation of more reduced compounds (Chakrawal et al., 2020; 2022; Kästner et al., 2024). However, the oxidation state of organic substrates can also affect microbial anabolism directly, because strongly oxidized molecules may require energy-costly reduction in order to be used for biomass synthesis (Von Stockar, 2010; Kästner et al., 2024). Links between substrate energetics and microbial carbon metabolism have been made across different scales in anaerobic systems (Keiluweit et al., 2016; Boye et al., 2017; Garayburu-Caruso et al., 2020a). However, it is yet unknown if energetic differences in substrate chemistry affect soil microbial carbon metabolism of aerated soils across pedo-climatic gradients in a predictable pattern.

In this study, we investigated whether and how bulk soil microbial carbon metabolism of aerated soils is linked to the carbon oxidation state and chemical composition of microbially available organic matter at the macroscale. We characterized KCl-extractable organic matter (as a proxy for microbially available) along a large geoclimatic gradient of temperate grassland soils using Fourier-transform ion cyclotron resonance mass spectrometry (LC-FT-ICR MS) and linked it to the measurements of specific growth and respiration as well as CUE collected in a previous study (Wasner et al., 2024b). We used regression models to explain the “full” range of growth, respiration and CUE, as well as the “residual” variation that was left unexplained in (Wasner et al., 2024b). Assuming that microbial carbon metabolism is energy-limited, we expected that microbial carbon turnover would be accelerated in soils where more oxidised organic carbon (i.e., higher NOSC) is available, because oxidation of such compounds can provide more energy per electron under aerobic conditions. At the same time, we expected microbial CUE to be lower in such soils, because more carbon needs to be “wasted” per electron harvested. We further examined whether we could find links between the distribution of different chemical classes of organic matter and microbial carbon metabolism along the geoclimatic gradient. We expected that the dominance of aliphatic compounds would favor higher rates of specific growth and respiration.

## 2. Material and Methods

### 2.1 Soil sampling and equilibration

We sampled 33 grassland A-horizon topsoils (0 - 10 cm) across a geoclimatic gradient in Chile in the summer seasons of 2017 and 2018. To increase the identifiability of the controls on microbial turnover, the gradient was constrained to a coherent type of land use and organic matter input (natural grassland and extensive rangeland) as well as carbonate free conditions. The gradient is described and characterized in full detail in previous studies (Doetterl et al., 2015; Wasner et al., 2024b; 2024a). Briefly, the gradient covers a range of water balance (MAP - PET) from 1704 to -1207 mm, and a MAT range from 3.0 to 17.1°C. The Köppen-Geiger climate zones range from arid steppe (Bsk) in the north to polar tundra (ET) in the south, excluding climatic extremes (hot and cold desert environments). Most sites (28) are situated in the temperate climate zone, representing climates with cold and warm summers, with or without dry season (Cfb, Cfc, Csb, Csc). Total soil organic carbon content spans from 6 to 188 g kg^-1^, and microbial biomass carbon from 7 to 464 μg C g^-1^ soil. After collection, soil samples were frozen at field moisture at -20°C and were stored and shipped in this condition. In the laboratory in Switzerland, samples were thawed, sieved to <2mm, and frozen again at -20°C until further analysis. Before all measurements (i.e., soil microbial carbon metabolism as well as organic matter extraction), soils were incubated for one week at 50% water holding capacity (WHC) and 20°C. Thereby, direct temperature and moisture limitation of microbial activity was eliminated. Measured microbial growth and respiration rates consequently reflect potential rates, and not necessarily *in situ* rates. While absolute values may deviate from *in situ* conditions, the patterns along the gradient nevertheless allow for mechanistic interpretation.

### 2.2 Measurement of microbial growth, respiration and CUE

The data describing soil microbial carbon metabolism along the gradient was previously published in (Wasner et al., 2024b) and methods are described in full detail there. Briefly, microbial growth rates and CUE were determined through ^18^O-H_2_O incorporation into DNA following a modified protocol based on (Spohn et al., 2016). Such growth estimates can serve as a proxy for parts of microbial anabolism (excluding extracellular polymeric substances and enzymes, (Geyer et al., 2019)). Microbial respiration rates were quantified with a LI-840A infrared gas analyzer (Li-Cor Inc., Lincoln, United States) in a continuous flow system including the Flux Puppy app (v1.0.0) (Carbone et al., 2019). Heterotrophic soil respiration can serve as a proxy for parts of aerobic catabolism, the process by which microbes extract energy from organic substrates (excluding catabolic processes such as e.g. fermentation to dissipate entropy, (Von Stockar, 2010)). Microbial biomass carbon (MBC) was quantified with chloroform fumigation extraction (modified from (Vance, Brookes and Jenkinson, 1987)). Microbial growth and respiration rates were expressed as specific rates per unit biomass (% of MBC day^-1^). Specific growth rates can serve as a community-weighted measure for the rate of microbial biomass turnover, assuming a stable amount of microbial biomass during incubation. Specific respiration rates are a community-weighted measure for aerobic catabolic activity per biomass. Specific growth rates ranged from 1 to 11 % of MBC day^-1^, specific respiration rates from 3 to 68 % of MBC day^-1^ and CUE from 0.1 to 0.6.

### 2.3 Chemical characterization of extractable organic matter using LC-FT-ICR MS

We characterized the chemical composition of organic matter that is potentially available to soil microbes using reversed phase liquid chromatography coupled to Fourier transform ion cyclotron resonance mass spectrometry (LC-FT-ICR MS). We extracted 7g of soil (at 50% WHC) for 1h with 35ml of 1M KCl on a horizontal shaker. The extracts were centrifuged (Sigma 3-16KL, 1730 x g, 15min, 4°C) and syringe-filtered to 0.45µm (Chromafil Xtra CA45/25, Macherey Nagel, pre-rinsed with 1M KCl) and subsequently stored at 4°C. To avoid organic contamination, all glassware was muffled (5h at 450°C), and all plastic material was acid washed with 0.1M HCl and rinsed with ultrapure water. Organic carbon concentration of the extracts was determined with a TOC/TN analyzer (TOC-L CPH/CPN, Shimadzu) and further used without subsequent solid-phase extraction. This was done to retain highly polar organic compounds. LC-FT-ICR MS was performed at the Helmholtz-Centre for Environmental Research in Leipzig (Germany), following the procedure described in (Lechtenfeld et al., 2024). Briefly, 100 µl of each acidified KCl extract were separated on a reversed-phase UHPLC column (ACQUITY UPLC HSS T3 VanGuard, 100 Å, 1.8 μm, 2.1 mm × 5 mm) using a water/methanol gradient adjusted with NH_4_OH (0.1 % formic acid adjusted with ammonia) to pH3. The injection order (also of the triplicated samples) was randomized, and quality control samples were added. The column outlet was directly connected to a 12T FT-ICR MS (SolariX xR, Bruker Daltonics, Billerica, USA) and the eluting DOM was ionized using an electrospray ionization source (Apollo II, Bruker Daltonics) operated in negative mode (4.2 kV capillary voltage). The ion accumulation time (IAT) was set to 0.8 s, and the mass range was set to m/z 147−1000. The mass resolving power (m/Δm, full width half-maximum) at m/z 400 was approximately 584,000 ± 14,000.

Chromatograms were segmented into twelve 1-min bins between 12 and 24 min retention time according to (Han et al., 2021), internally recalibrated using known DOM masses, and molecular formulas assigned to mass peaks > S/N 4 with the Lambda-Miner (Wurz et al., 2024) including elements C (0-60), H (1-122), N (0-2), O (0-40), and S (0-1). Molecular formulas in process and instrumental blanks (*n =* 8) were removed from the data and retention time segments flattened into a pseudo-DI spectrum using the mean of raw intensities.

For one soil of low, intermediate, and high SOC content (11, 63 and 151 g SOC kg ^-1^, respectively) the measurements were made in triplicate, starting from pre-incubation and following through the entire workflow. Visual analysis shows that the chosen methods are very reproducible and sensitive enough to characterize the distinct composition of extractable organic matter along the gradient (Figure S7).

Due to direct injection of extracted carbon into LC-FT-ICR MS without dilution, the intensities of 30.5 % of the molecular formulas correlated (false discovery corrected q-value < 0.05) with the organic carbon concentration in the extracts (ranging from 0.8 to 12.7 mg C L^-1^) (Figure S2). To account for this effect and thereby obtain a measure for the “relative” composition of extractable organic matter, we normalized all peak intensities to the organic carbon concentration of the respective sample.

As a proxy for the carbon oxidation state of extracted organic matter we estimated the nominal oxidation state of carbon (NOSC) for each molecular formula. We based the estimation of NOSC on molecular stoichiometry, following the empirical formula from (LaRowe and Van Cappellen, 2011):

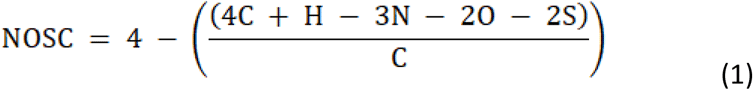

where the letters indicate the number of each chemical element in a molecular formula. We assigned the molecular formulas to chemical classes based on O/C and H/C ratios and a modified aromaticity index (AI_mod_). AI_mod_ was calculated based on (Koch and Dittmar, 2016):

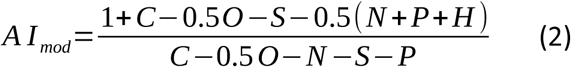

where the letters indicate the number of each chemical element in a molecular formula. Formulas with AI_mod_ > 0.5 *mus*t contain an aromatic ring. The chemical classification into five broad classes followed (Hawkes et al., 2020), where “aliphatic” was defined as H/C ≥ 1.5, “low O unsaturated” as H/C < 1.5 and AI_mod_ ≤ 0.5 and O/C < 0.5, “high O unsaturated” as H/C < 1.5, AI_mod_ ≤ 0.5 and O/C ≥ 0.5, “aromatics” as 0.5 < AI_mod_ < 0.67, and “condensed aromatics” as AI_mod_ ≥ 0.67. We chose this classification scheme because it is conservative and leaves no gaps with unspecified compounds which facilitates the discussion of results. Given the uncertainty that is inherent in chemical classification based on element ratios, chemical class names reflect similarity to the respective classes rather than confirmed membership (Laszakovits and MacKay, 2022).

### 2.4 Statistical evaluation of the mass spectra data

To identify outliers potentially caused by sample contamination, we plotted the total signal intensity of all formulas against (i) the number of formulas and against (ii) organic carbon concentration of the extracts per sample (Figure S1a,c). Four samples were visually found to contain outlier formulas. We therefore filtered the entire dataset for peaks (i) with intensities larger than the 95th percentile across all samples and peaks, which (ii) were outliers following the interquartile rule, and (iii) were present in four or less samples. The intensities of these outlier peaks were then imputed by *k*-nearest neighbors estimation based on the mass spectra of the most similar other samples (R-package “DMwR2”; (Torgo, 2016)) (Figure S1b,d).

We applied regression statistics to investigate links between substrate chemistry and bulk microbial carbon metabolism. The six response variables that reflect bulk microbial carbon metabolism were the specific growth and respiration rates and carbon use efficiency, as well as the residuals that were left for these variables after stepwise regression in a previous study (Wasner et al., 2024b).

We applied simple linear regression to test whether the six response variables are linked to intensity-weighted mean NOSC across the gradient. Residuals were tested for normal distribution (Shapiro-Wilk test) and homoscedasticity (Breusch-Pagan test, R-package “lmtest”, (Zeileis and Hothorn, 2002)). Where either of the two was not fulfilled, the response variables were transformed with the natural logarithm.

We applied cross-validated partial least squares (PLS) regression to test if and how relative substrate composition (i.e., the normalized peak intensities) is linked to the six response variables. We scaled the normalized peak intensities, and split the dataset 100 times into a 80 % training set and a 20 % validation set. To avoid overfitting, we chose the best model based on the RMSE of the predictions of the left-out validation data averaged over all 100 iterations, and we limited the maximum possible number of latent variables to ten. PLS regression was performed with the R-package “caret” (Kuhn, 2008) which wraps the package “pls” (Liland, Mevik and Wehrens, 2024). PLS regression requires a choice: On the one hand, PLS regression fails (and computational cost is prohibitive) if the independent data is too sparse and features too many variables. On the other hand, removing too many variables prior to regression can potentially result in information loss. As a trade-off, we chose to only consider molecular formulas that occurred in at least half (>17) of the soil samples for the predictor dataset (i.e., 5305 formulas). Sensitivity analysis shows that the interpretation of the results was not affected by this choice (Figure S3).

### 2.5 Interpretation of PLS regression results

The amount of variability explained by a PLS model can be described with the R ^2^ value. However, note that PLS regression does not compute p-values for the coefficients. Following PLS regression, we used the varImp() function from “caret” to identify the top 10% most important chemical formulas that featured as predictors in each model. This threshold represents another choice, where consideration of too many or too few predictor variables can potentially result in information loss. Sensitivity analysis shows that interpretation of the results was not affected by this choice (Figure S4).

PLS regression identifies the molecular formulas that best align with the dependent variable. Since spectral data inherently exhibits autocorrelation (Figure S5), one cannot fully rule out that some selected compounds were associated due to autocorrelation rather than causal relationship. However, supplementary correlation analysis showed that correlation among the top 10% most important predictive formulas (Figure S6) was generally low to moderate.

To assess which chemical classes were particularly important in the models, we calculated an importance factor (IF) for each chemical class in each model. This is necessary, because molecular formulas from chemical classes that are more numerous in the dataset than others (e.g., high O unsaturated > condensed aromatics, Table 1) have a higher likelihood to feature among the top 10% predictive molecular formulas, thereby confounding the identification of important chemical classes. The IF was calculated for each chemical class in each model:

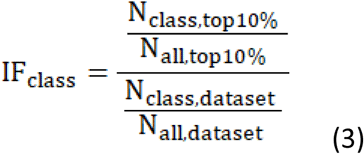

where *N* is the number of chemical formulas, the subscript *class* refers to the specific chemical classes, the subscript *all* refers to all chemical classes, the subscript *top10%* refers to the top 10% predictive molecular formulas, and the subscript *dataset* refers to the entire predictor dataset. An IF value >1 indicates that a chemical class is overrepresented in the top 10% predictive molecular formulas as compared to the predictor dataset, and therefore likely particularly relevant for the response variable, while an IF value <1 means the opposite.

**Table 1.**
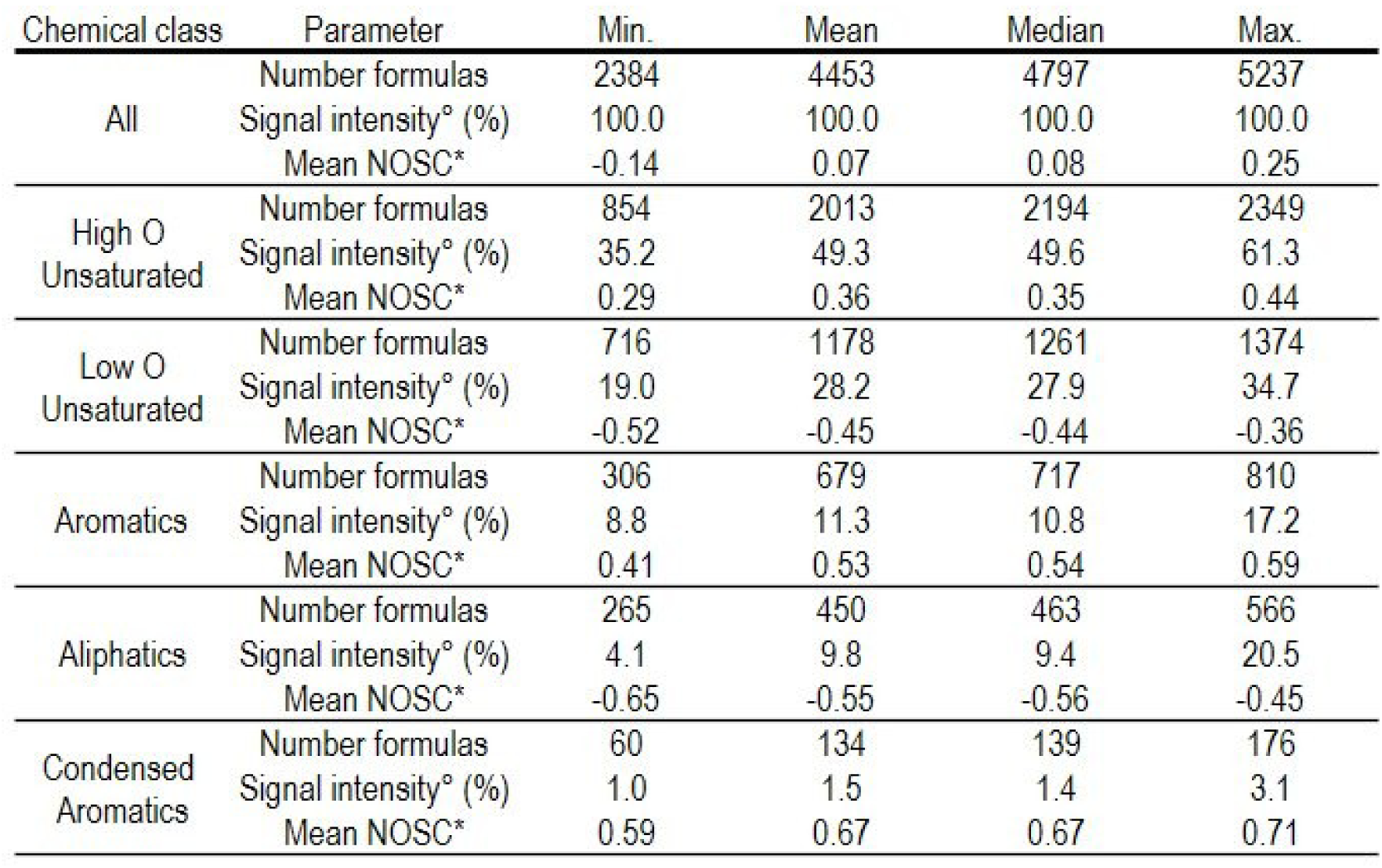
Summary of the chemical composition of extractable organic matter. Minimum (min.), mean, median and maximum (max.) values are shown across the 33 investigated soils, for all chemical classes together and for the individual chemical classes. Shown is data for the subset of molecular formulas which were used for PLS regression (see method section 2.4). A summary of the entire mass spectra (i.e., including rare molecular formulas not considered for PLS regression) is shown in Table S1. °Percent of summed signal intensities per sample; *Intensity-weighted mean. NOSC = Nominal oxidation state of carbon.

| Chemical class | Parameter | Min. | Mean | Median | Max. |
| --- | --- | --- | --- | --- | --- |
| All | Number formulas | 2384 | 4453 | 4797 | 5237 |
|  | Signal intensity<sup>o</sup> (%) | 100.0 | 100.0 | 100.0 | 100.0 |
|  | Mean NOSC* | -0.14 | 0.07 | 0.08 | 0.25 |
| High O<br>Unsaturated | Number formulas | 854 | 2013 | 2194 | 2349 |
|  | Signal intensity<sup>o</sup> (%) | 35.2 | 49.3 | 49.6 | 61.3 |
|  | Mean NOSC* | 0.29 | 0.36 | 0.35 | 0.44 |
| Low O<br>Unsaturated | Number formulas | 716 | 1178 | 1261 | 1374 |
|  | Signal intensity<sup>o</sup> (%) | 19.0 | 28.2 | 27.9 | 34.7 |
|  | Mean NOSC* | -0.52 | -0.45 | -0.44 | -0.36 |
| Aromatics | Number formulas | 306 | 679 | 717 | 810 |
|  | Signal intensity<sup>o</sup> (%) | 8.8 | 11.3 | 10.8 | 17.2 |
|  | Mean NOSC* | 0.41 | 0.53 | 0.54 | 0.59 |
| Aliphatics | Number formulas | 265 | 450 | 463 | 566 |
|  | Signal intensity<sup>o</sup> (%) | 4.1 | 9.8 | 9.4 | 20.5 |
|  | Mean NOSC* | -0.65 | -0.55 | -0.56 | -0.45 |
| Condensed<br>Aromatics | Number formulas | 60 | 134 | 139 | 176 |
|  | Signal intensity<sup>o</sup> (%) | 1.0 | 1.5 | 1.4 | 3.1 |
|  | Mean NOSC* | 0.59 | 0.67 | 0.67 | 0.71 |

### 2.6 Redundancy analysis

We performed redundancy analysis (RDA) to explore potential relationships between organic matter chemistry and potential environmental drivers. We applied Hellinger-transformation on the normalized peak intensities of the mass spectral subset which was used for PLS regression. As potential predictor variables we used two datasets: rotated components describing environmental conditions (based on climate and soil physicochemistry) and rotated components describing microbial community composition (based on relative abundances of dominant bacterial and fungal genera). Data was taken from (Wasner et al., 2024b), and underlying methods are described in detail there. We performed forward selection with 999 permutations (R-package “vegan”, (Oksanen et al., 2025)) to identify significant predictors (p-value < 0.05) from each of the two datasets, and applied permutation based ANOVA to assess the significance of the resulting RDA models.

## 3. Results

### 3.1 Chemistry of extractable organic matter along the gradient

The extracted and filtered (< 0.45µm) organic matter accounted on average for 0.24 % of total organic carbon (0.05 to 0.66 %). This amount was in the expected range (Gao et al., 2024) and was in the same order of magnitude as the amount of carbon which was taken up (i.e., growth + respiration) by the microbial communities over 24h (on average 58 % of uptake, ranging from 15 to 177 %). The average redox status (i.e., intensity-weighted mean NOSC) of the extractable organic matter varied between -0.14 to 0.25 across the gradient. Although direct cross-study comparisons of intensity-weighted mean NOSC are difficult due to different extraction and mass spectrometry methods, the values in this study were similar to previous values reported for water extractable organic matter measured in positive mode (-0.47 to -0.16; (Dufour et al., 2022); around -0.06; (Simon et al., 2025)). Classification of molecular formulas into chemical classes following (Hawkes et al., 2020) allowed to characterize the composition of extractable organic matter. The most numerous chemical classes across the entire gradient were unsaturated compounds, with high as well as low O/C ratios, contributing between 35 to 61 % to total signal intensities (Table 1). Aliphatics, aromatics and condensed aromatics each contributed only between 1 to 21 % to total signal intensities (Table 1). RDA revealed that variation in organic matter composition was partly related to environmental conditions and microbial community composition (25.3 and 40.4 % of variation explained in two independent RDA analyses, Table S3).

### 3.2 Links between chemistry of extractable organic matter and microbial functions

The intensity-weighted mean NOSC explained only a small amount of variation of “full” CUE (p-value = 0.03, adj. R^2^ = 0.12, coefficient = 1.15), and none for the other five response variables (Table S2). In contrast, the chemical composition of extractable organic matter explained more than half of the variation (R^2^ between 0.57 and 0.67 for the “full” models). Substrate chemistry was partly redundant to the environmental factors investigated in a previous study (namely microbial community composition, soil physicochemistry and climate; (Wasner et al., 2024b)), as indicated by the lower R^2^ values for the “residual” models (0.27 to 0.43, Figure 2d-f) compared to the “full” models (0.57 to 0.67, Figure 2a-c). The molecular formulas which explained full specific growth and full specific respiration rates were similar in the region of higher H/C ratios, and different in the region of lower H/C ratios (Figure 2a,b). In contrast, full CUE was most strongly related to a different set of molecular formulas with low H/C ratios and high O/C ratios (Figure 2c). The molecular formulas linked to residual variation of specific growth and respiration rates (Figure 2d,e) differed from those linked to full rates (Figure 2a,b), with a shift to lower H/C ratios. The model for residual CUE (Figure 2f) shifted to negative relationships with molecular formulas of even higher O/C ratios, as compared to the model for full CUE (Figure 2c).

**Figure 1.**
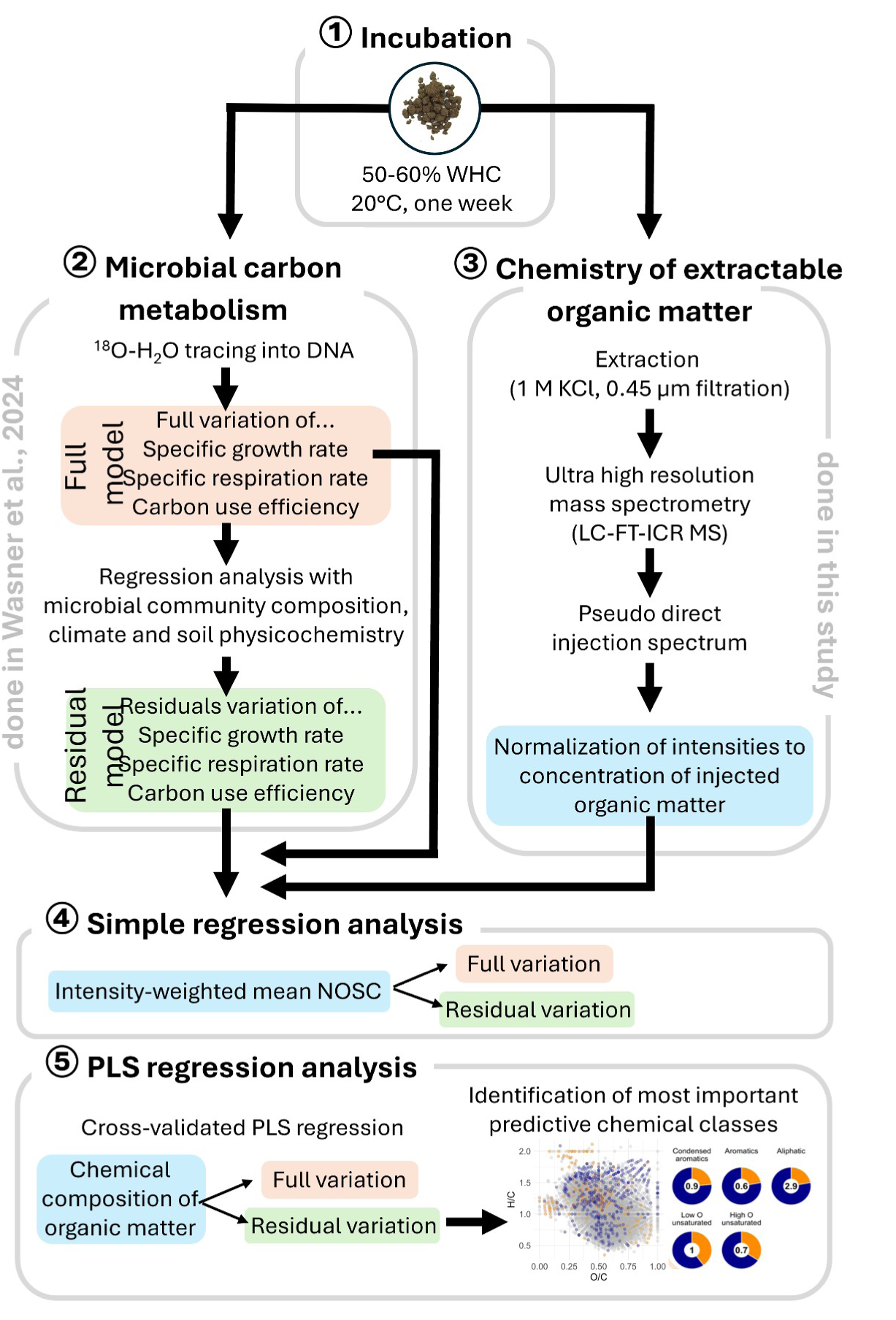
Summary of workflow. ① Soils were incubated under standardized conditions. ② In a previous study (Wasner et al., 2024b), bulk microbial carbon metabolism were measured via ^18^O-H_2_O labelling, and regression analysis was applied to identify the drivers of bulk microbial carbon metabolism (microbial community composition, climate and soil physicochemistry). ③ The chemical composition of extractable organic matter was determined using reversed phase liquid chromatography in combination with Fourier transform ion cyclotron resonance mass spectrometry (LC-FT-ICR MS). ④ Lastly, regression analysis was conducted to identify chemical classes which are linked to the investigated bulk microbial carbon metabolism.

**Figure 2.**
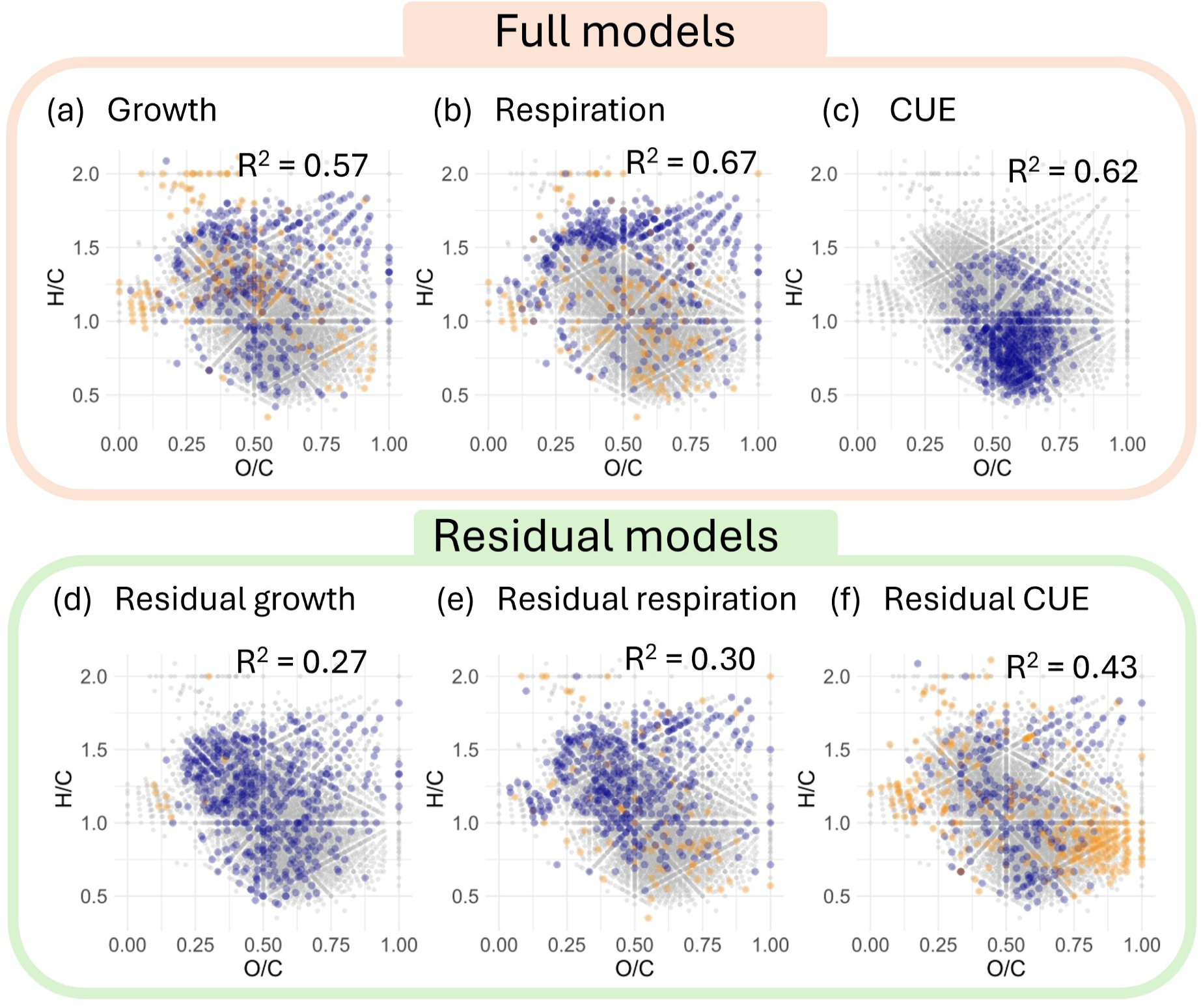
Van Krevelen diagrams showing the most important predictive molecular formulas for each of the six models. Shown in grey are all molecular formulas, highlighted in dark blue are the top 10% most important formulas with positive coefficients, and highlighted in light-orange are the top 10% most important formulas with negative coefficients. The upper panels (a) to (c) show the “full” models where models were fitted to the full variation of growth, respiration and carbon use efficiency (CUE), while the lower panels (d) to (f) show the “residual” models where models were only fitted to the residuals from (Wasner et al., 2024b).

Classification of molecular formulas into chemical classes enabled us to identify which types of compounds the microbial functions were linked to. Full specific growth and respiration rates (Figure 3a,b) were most strongly positively linked to aliphatic compounds. Full specific growth rates also had a predominantly negative link to compounds with high H/C and low O/C ratios (upper left corner of van Krevelen diagram), to which full respiration rates were predominantly positively linked (Figure 2a,b). Full CUE was mainly positively linked to aromatics and condensed aromatics (Figure 3c). Considering only residual variation, specific growth and respirations rates were mainly positively linked to aliphatic compounds and unsaturated compounds with low O (Figure 3d,e). Residual CUE was most strongly linked to aliphatic compounds, with mixed coefficients (Figure 3f). Except for the models for CUE and residual growth, the degree of autocorrelation among the top 10% most important predictive molecular formulas was generally moderate to low (Figure S6). This means that the model structures are unlikely to be solely a product of autocorrelation among molecular formulas.

**Figure 3.**
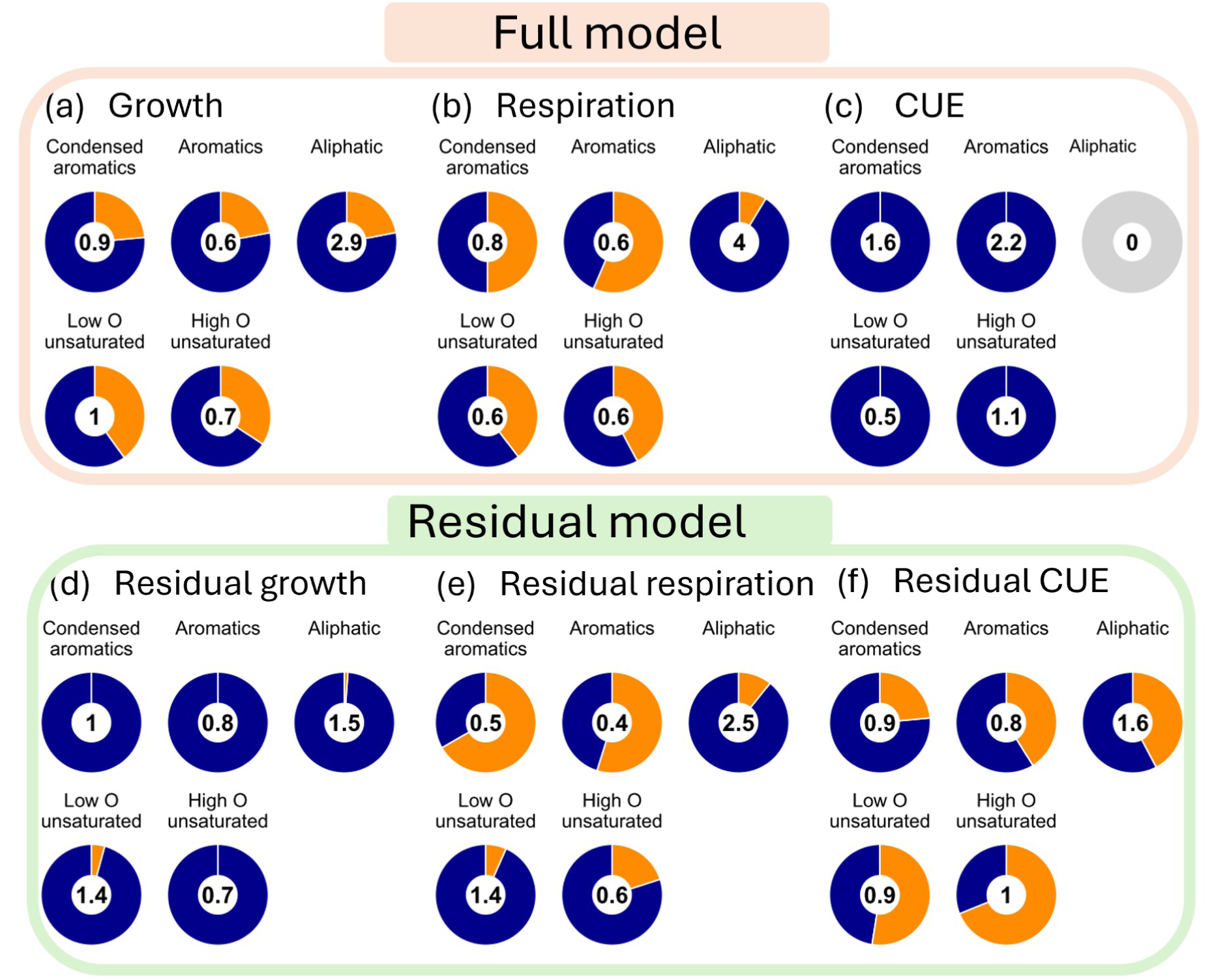
Contribution of the assigned chemical classes to the top 10% most important predictive molecular formulas for each of the six models. The numbers show the relative importance (IF, see method section 2.5) of each chemical class for the respective model; values >1 indicate high relevance, values <1 low relevance. Color indicates positive (dark-blue) versus negative (light-orange) coefficients. The upper panels (a) to (c) show the “full” models where the full variation of growth, respiration and carbon use efficiency (CUE) was fitted, while the lower panels (d) to (f) show the “residual” models where only the residuals from (Wasner et al., 2024b) were fitted.

Full growth was positively linked to similar numbers of molecular formulas with NOSC larger and smaller than 0, but most negative links were with formulas with negative NOSC (Figure 4a). In contrast, molecular formulas that were positively linked to full respiration mostly had negative NOSC (Figure 4b). The same patterns were even more pronounced for residual variation of growth and respiration rates (Figure 4d,e). Full CUE was almost entirely linked to molecular formulas with positive NOSC (Figure 4c). In contrast, residual CUE showed positive and negative links across molecular formulas with positive and negative NOSC, with the strongest negative associations to molecular formulas of even higher NOSC between 0.5 to 1 (Figure 4f).

**Figure 4.**
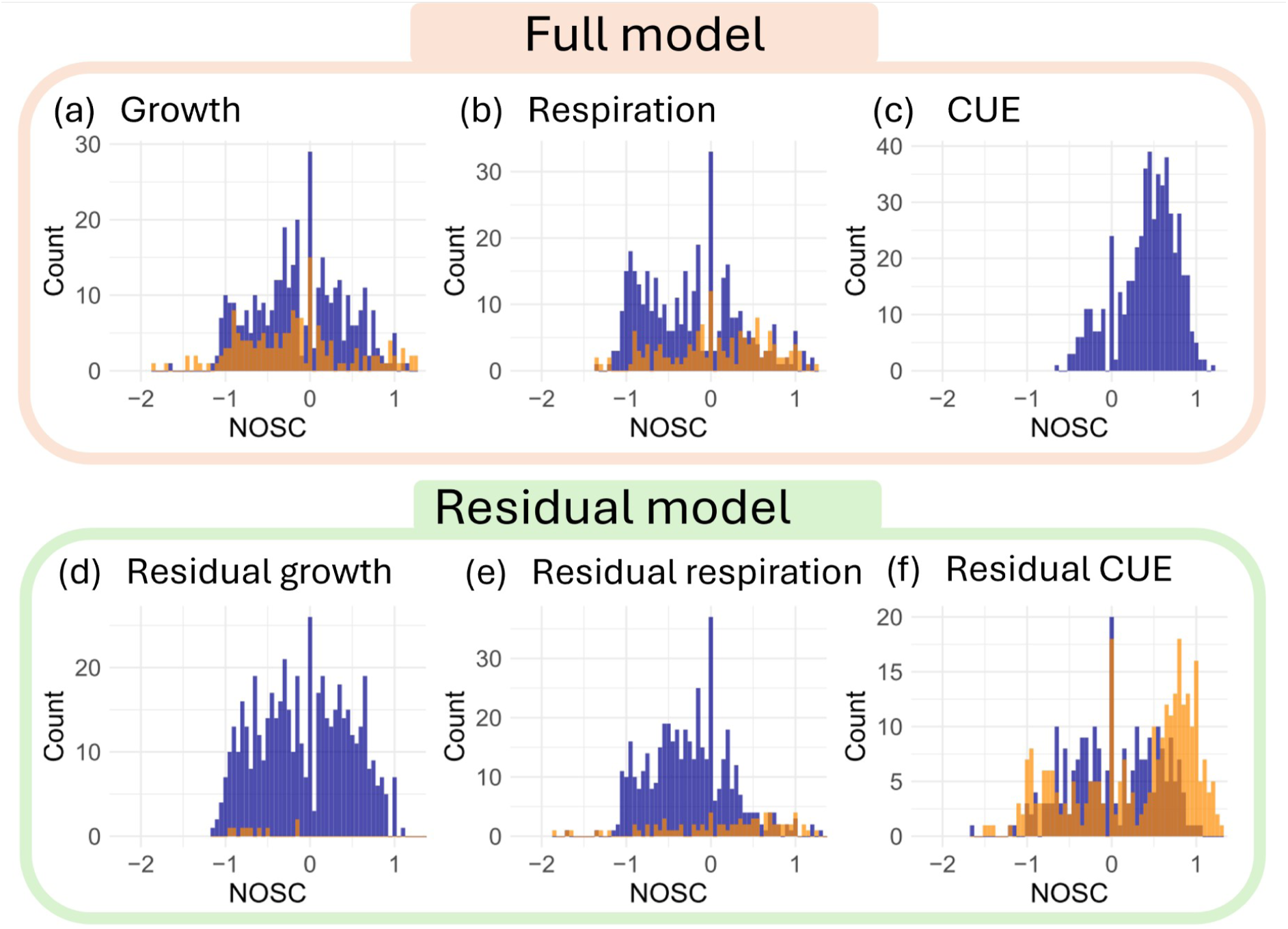
Distribution of NOSC across the top 10% most important predictive molecular formulas for each of the six models. Color indicates positive (dark-blue) versus negative (light-orange) coefficients. The upper panels (a) to (c) show “full” models where the full variation of growth, respiration and carbon use efficiency (CUE) was fitted, while the lower panels (d) to (f) show “residual” models where only the residuals from (Wasner et al., 2024b) were fitted. The bin width is 0.05 NOSC.

## 4. Discussion

### 4.1 Mean oxidation state of carbon does not explain bulk microbial carbon metabolism

To test our first hypothesis —that more oxidized substrates lead to faster microbial carbon turnover and lower CUE— we calculated the intensity-weighted mean NOSC for each soil to simplify the complex mass spectrometry data. However, the intensity-weighted mean NOSC barely explained any variation in specific growth and respiration rates or CUE (Table S2). This lack of a relationship could indicate that microbial carbon metabolism is unrelated to substrate NOSC under aerobic conditions. However, it is also possible that the intensity-weighted mean NOSC is an unsuitable oversimplification that is not necessarily biologically meaningful. The extractable organic matter was dominated by unsaturated compounds (Table 1). This range of the van Krevelen diagram has been consistently associated with lignin- and tannin-like compounds (Boye et al., 2017; Laszakovits and MacKay, 2022), which have also dominated extractable organic matter in other studies (Freeman et al., 2024; Shi et al., 2025). However, while these chemical classes were dominant and affected intensity-weighted mean NOSC (Table 1), they were not proportionally linked to microbial carbon turnover (Figure 3). Instead, extractable organic matter can be enriched with lignin- and tannin-like compounds because they are abundant in plant organic matter input and/or because microbes are thought to preferentially decompose and transform other compounds (Kraus, Dahlgren and Zasoski, 2003; Lützow et al., 2006; Grandy and Neff, 2008; Jones et al., 2023). Overall, our finding emphasizes that weighted averages are not sensitive enough to be linked to microbial metabolism. Simplified intensity-weighted mean NOSC is not necessarily biologically meaningful because it also includes compounds that may not be used by microbes. Rather, molecular data is required in which we need to identify the (few) indicative marker formulas that can be linked to metabolism (e.g. as in Stumpf et al., 2025).

### 4.2 Bulk microbial carbon metabolism is linked to the chemical composition of extractable organic matter

More than half of the variation of specific growth (R^2^ = 0.57), specific respiration (R^2^ = 0.67) rates and CUE (R^2^ = 0.62) could be explained with the relative chemical composition of extractable organic matter (Figure 2). The strong links between extracted compounds and bulk microbial carbon metabolism indicate that parts of the KCl-extracted organic matter are microbially available substrate (otherwise there would be no links).

Substrate chemistry was partly redundant to the environmental factors investigated in a previous study (namely microbial community composition, soil physicochemistry and climate; Wasner et al., 2024b), as indicated by the lower R^2^ values for the “residual” models (0.27 to 0.43, Figure 2d-f) compared to the “full” models (0.57 to 0.67, Figure 2a-c). This redundancy was expected, given that organic matter chemistry is a complex product of climate and mineralogy, quantity and qualitative properties of plant inputs and microbial transformation of organic matter (Table S3, (Roth et al., 2015; Hall et al., 2020; Mainka et al., 2022)). Quantity and quality of organic matter may co-determine microbial community composition (Cederlund et al., 2014; Fierer, 2017; Wasner et al., 2025), which in turn affects substrate use (Trivedi, Anderson and Singh, 2013; Malik et al., 2020; Piton et al., 2023) and bulk microbial carbon metabolism (Graham et al., 2016; Wasner et al., 2024b). This way, the composition of the active microbial community may feed back to the chemical composition of organic matter (Domeignoz-Horta et al., 2021; Campbell et al., 2022; Flamholz et al., 2025). Despite these complex redundancies between environmental characteristics and organic matter chemistry (Table S3), the overall links between specific growth and respiration and organic matter chemistry remained consistent when we accounted for the redundancy by only predicting the residuals of variation (“residual” models) (Figure 3). The explained variation in the “residual” models therefore provides evidence that microbial metabolism is linked to the chemical composition of extractable organic matter beyond mere covariation. In the following section, we discuss the observed links in more detail.

### 4.3 Rates of catabolism and anabolism are positively linked to aliphatic and unsaturated compounds

Higher rates of microbial catabolism (i.e., specific respiration) and - to a slightly lesser degree - anabolism (i.e., specific growth) were linked to aliphatic compounds (Figure 3), which is in line with expectations from literature (Gunina and Kuzyakov, 2022). In soils in which aliphatic compounds were more dominant in the extractable organic matter pool, microbes grew and respired faster (as compared to soils where they were less dominant). Examples of aliphatic compounds associated with the discussed range in the van Krevelen diagram include carbohydrate-, peptide-/protein- as well as amino sugar-like molecules (Boye et al., 2017; Laszakovits and MacKay, 2022). All three chemical classes are abundant in soils, in part because they have a high affinity to become stabilized on mineral surfaces and thereby accumulate (Quiquampoix and Burns, 2007; Kögel-Knabner and Amelung, 2014; Kallenbach, Frey and Grandy, 2016). Carbohydrates are heterogenous, derived from plant material and microbial biomass, and are known to be a primary energy source for soil microbes (Gunina and Kuzyakov, 2015; Chantigny, Olk and Angers, 2025). Amino sugars mainly derive from microbial biomass, where they are structural compounds of cell walls (Chantigny, Olk and Angers, 2025). Proteinaceous compounds have a high N content due to their N-amide functional groups, which makes them a main source of organic N for soil microbes (Quiquampoix and Burns, 2007; Kögel-Knabner and Amelung, 2014).

In the “residual” models, specific growth and respiration were further linked to unsaturated compounds with low O (Figure 3d,e) which have elsewhere been associated with lignin-like and lignin-derived compounds (Boye et al., 2017; Laszakovits and MacKay, 2022). Although lignin decomposition is considered to be comparatively slow (Hall et al., 2020), lignins can be oxidized by microbes that possess the necessary enzymatic toolboxes, particularly fungi (Sinsabaugh, 2010; Atiwesh et al., 2022). In soils where enough oxidative enzymes are present, lignin degradation may therefore be a reasonable driver of soil microbial metabolism, secondary to more favorable aliphatic substrates. The fact that both - specific growth *and* respiration - were associated with aliphatics (such as carbohydrates, peptides/proteins and amino sugars) and unsaturated compounds (such as lignins) could indicate either that (i) these compounds are readily oxidised and the obtained energy is instantly used to fuel anabolism (in the case that soil microbes are energy limited), or (ii) that these compounds allow for more anabolism through direct incorporation into biomass, subsequently leading to higher catabolic rates (in the case that soil microbes are substrate limited). Additional experiments would be required to determine which type of limitation underlies the observed relationship.

### 4.4 Carbon use efficiency appears to be linked to oxidized compounds

Surprisingly and in contrast to our expectation, the full variation of CUE was almost exclusively positively linked with compounds classified as aromatics (defined as 0.5 < AI_mod_ < 0.67) and condensed aromatics (defined as AI_mod_ ≥ 0.67) (Figure 3c). Such compounds could be degradation products from lignins and tannins or may stem from incomplete combustion of biomass (Wilcke, 2000; Knicker, 2011; Bird et al., 2015). This relationship is unexpected for several reasons. (1) Highly aromatic compounds are comparatively oxidised (Table 1); in theory, this would mean that they require a low standard state Gibbs energy for the oxidation half reactions, and that they could therefore be energetically favorable for catabolism. If these compounds were primarily used for catabolism, this would lower CUE rather than increase it. (2) Aromatics and condensed aromatics do not feature among the most important predictive compounds for catabolism (Figure 3b). (3) To drive efficient carbon metabolism, the compounds would need to be incorporated into anabolism at low energetic cost, which is unlikely because they would need to be reduced beforehand. A possible explanation for this unexpected observation is that the effect of other chemical classes that drive specific growth and respiration cancels out, and that fungal metabolism tips the scales in this regression model. Fungi can produce oxidases and peroxidases which allow them to process and consume some types of aromatic compounds (Sinsabaugh, 2010; Khatoon et al., 2017; Deng et al., 2021), and it has been suggested that fungi might have a higher CUE than bacteria (He et al., 2024). Consequently, soils with higher CUE may have a comparatively higher share of active fungal metabolism, fueled by the abundance of certain aromatic compounds. When we only considered residual variation, this relationship disappeared entirely (Figure 3c,f), indicating that it was indeed redundant to aspects of microbial community composition that we found in (Wasner et al., 2024b). Instead, the residual variation in CUE showed a strong negative link with certain aliphatics and even more oxidised compounds (Figure 4) that were neither linked to growth nor respiration (Figure 2). The results indicate that these compounds render carbon use less efficient than would be expected based on environmental factors and microbial community composition alone. However, the identity of these compounds as well as the underlying mechanism require further research.

### 4.5 The oxidation state of carbon can explain catabolism but not anabolism

A main motivation of this study was to find out if bulk microbial metabolism of upland soils is linked to the carbon oxidation state of the microbial substrate at the macroscale. We observed that respiration was linked more strongly to reduced compounds (NOSC <0) in comparison to growth, which was positively linked to a range of compounds from reduced (NOSC = -1) to oxidized (NOSC = 1) (Figure 4). Furthermore, specific respiration rates were positively linked to aliphatics in the van Krevelen range that has been associated with lipid-like compounds (Figure 2, (Boye et al., 2017; Laszakovits and MacKay, 2022)). Lipids are a very broad chemical class that derives from plant material and microbial biomass (Kögel-Knabner and Amelung, 2014) and that is comparatively reduced (Wang and Kuzyakov, 2023). Similarly, we visually observed a predominantly positive link between specific respiration rates and unsaturated compounds with O/C < 0.25, possibly cutan or suberan (Supplementary discussion S1). Assuming causality, these observations could indicate that reduced (as compared to oxidised) compounds are preferably respired to obtain energy in soils where microbial carbon turnover is carbon-limited. A reason for preferential oxidation compounds with reduced carbon could be carbon conservation, because while oxidation of such compounds yields comparatively less energy per mole electron, more electrons can be removed per mole carbon and thus less carbon is wasted per unit energy (Chakrawal et al., 2022; Gunina and Kuzyakov, 2022).

In contrast, growth could be more disconnected from the carbon oxidation state of utilized compounds for several reasons. First, energy stored on electron carriers (obtained from catabolism) could be used to facilitate energetically unfavorable transformations (i.e., reduction) of compounds that are required for anabolism (Wang and Kuzyakov, 2023). Second, mining of “building blocks” that directly enter anabolism through salvage pathways could allow growth based on a wider range of compounds irrespective of their carbon oxidation state ((Gunina and Kuzyakov, 2022) and references therein; (Kästner et al., 2021; Wasner et al., 2023)). Salvage pathways could consequently partly decouple anabolism from catabolism through the direct use of compounds in anabolism without large energy investments into chemical transformation (Kästner et al., 2021).

Another observation that supports the conclusion that the carbon oxidation state of substrate may not be the single most important substrate feature for microbial growth is that across and within chemical classes, NOSC did not determine whether a compound was positively or negatively related (across classes: Figure 4; within classes: not shown). In contrast, compounds that were linked to microbial metabolism spanned a wide range of NOSC values (Figure 4), which corresponds with findings from a time-resolved metabolic footprinting study on a soil isolate (Cyle et al., 2020). Very likely, other aspects than carbon oxidation state also affect microbial substrate use, such as energy demand for metabolic machinery (Amenabar et al., 2017), salvage pathways (Kästner et al., 2021) or stoichiometric demands like the C:N ratio (Mooshammer et al., 2014; Chakrawal et al., 2022). Compound concentration and accessibility may also be dominant determinants of investments into metabolic machinery and microbial reaction kinetics, outweighing purely thermodynamic controls (LaRowe and Van Cappellen, 2011; Garayburu-Caruso et al., 2020b; Lehmann et al., 2020; Wang and Kuzyakov, 2023).

Overall, our findings suggest that catabolic activity may be more strongly linked to the carbon oxidation state of the substrate than microbial growth. Without further information about active metabolic pathways, substrate NOSC showed little relationship with specific growth rates. We argue that the links between bulk microbial metabolism and the chemical composition of extractable organic matter were likely related to other differences between the chemical classes, such as microbial toolboxes for their use, their concentrations or stoichiometric demands.

## 5. Conclusion

This study provides empirical evidence that rates of microbial catabolism and anabolism are linked to the chemical composition of organic matter, independent of microbial community composition and other environmental factors. In particular, we found that soils had a faster microbial carbon turnover per unit microbial biomass where aliphatic compounds (such as carbohydrate-, proteinaceous- and amino sugar-like) dominated extractable organic matter. A secondary source of rapid microbial carbon turnover seem to be unsaturated compounds, potentially lignin-like. Specific respiration rates were positively linked to reduced compounds, suggesting carbon-conserving catabolism. In contrast, links between specific growth rates and substrates could not be explained with the carbon oxidation state of the compounds. This indicates that even though soil microbes are widely assumed to be energy-limited, redox chemistry is not the critical lever to understand microbial growth in aerated upland soils. Rather, the links between organic matter chemistry and microbial carbon metabolism in such soils must be resolved through improved understanding of active biochemical pathways. Possibly, a promising next step could be to try and link metabolomic data from ultra high resolution mass spectrometry with metagenomic or metatranscriptomic data and growth rate estimates based on stable isotope probing, in order to identify biochemical pathways that are predictive of microbial carbon turnover rates. Our findings imply that chemical characterization of extractable organic matter can provide a way forward in the challenging and pressing task to understand, predict and upscale soil microbial carbon turnover.

## Acknowledgements

The authors would like to thank Erick Zagal Venegas for providing the soil samples and René Köhler for support with the laboratory work. D.W. would like to thank Louis Dufour for interesting exchanges about the topic.

## Supplementary discussion S1

We observed a cluster of unsaturated compounds with low O which were predominantly positively linked to specific respiration but negatively or not linked to specific growth. A common example for unsaturated hydrocarbons are alkenes, however those are thought to be comparatively easy to metabolize. Possibly, the unsaturated hydrocarbon compounds in question could stem from cutan and suberan, which can derive from the decomposition of abundant lipid plant biopolymers such as cutin (primarily leaf-derived) and suberin (primarily root-derived), and are hard to decompose for microbes (Kögel-Knabner and Amelung, 2014). While cutan and suberan are in principal saturated, it has been found that they can have minor contributions of unsaturated structures (Turner, Hartman and Hatcher, 2013; Leide et al., 2020) which could place them in the range of unsaturated compounds with very low O. The association of cutan and suberan with the cluster of unsaturated compounds with very low O is speculative, but future MS/MS experiments might help to elucidate the links between metabolism and potential suberin/cutin substructures.

---

**Figure S1.**
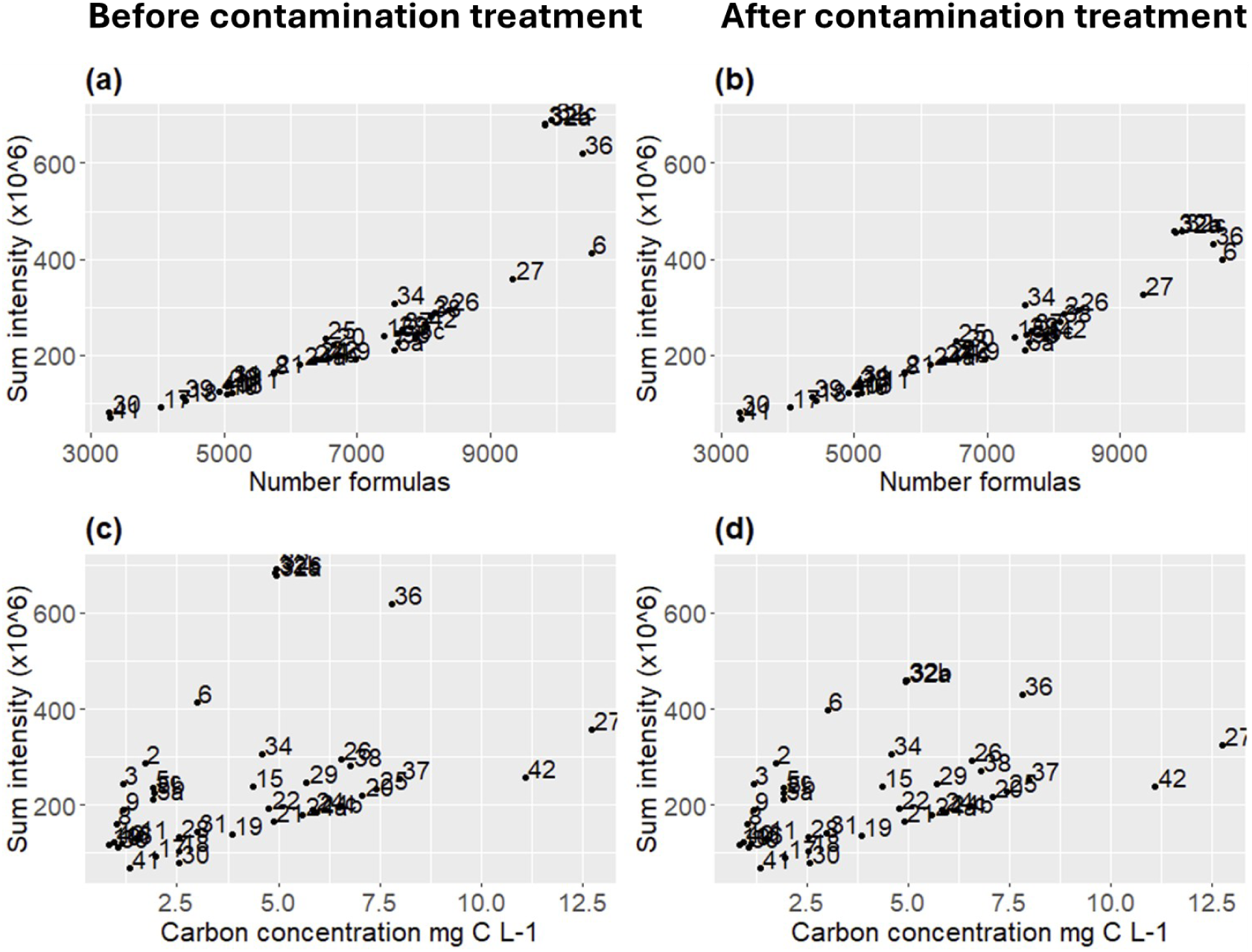
Total signal intensity before and after outlier treatment. Panels (a) and (b) show total signal intensity against the number of formulas per sample, and panels (c) and (d) against organic carbon concentration. Numbers indicate the sample ID.

**Figure S2.**
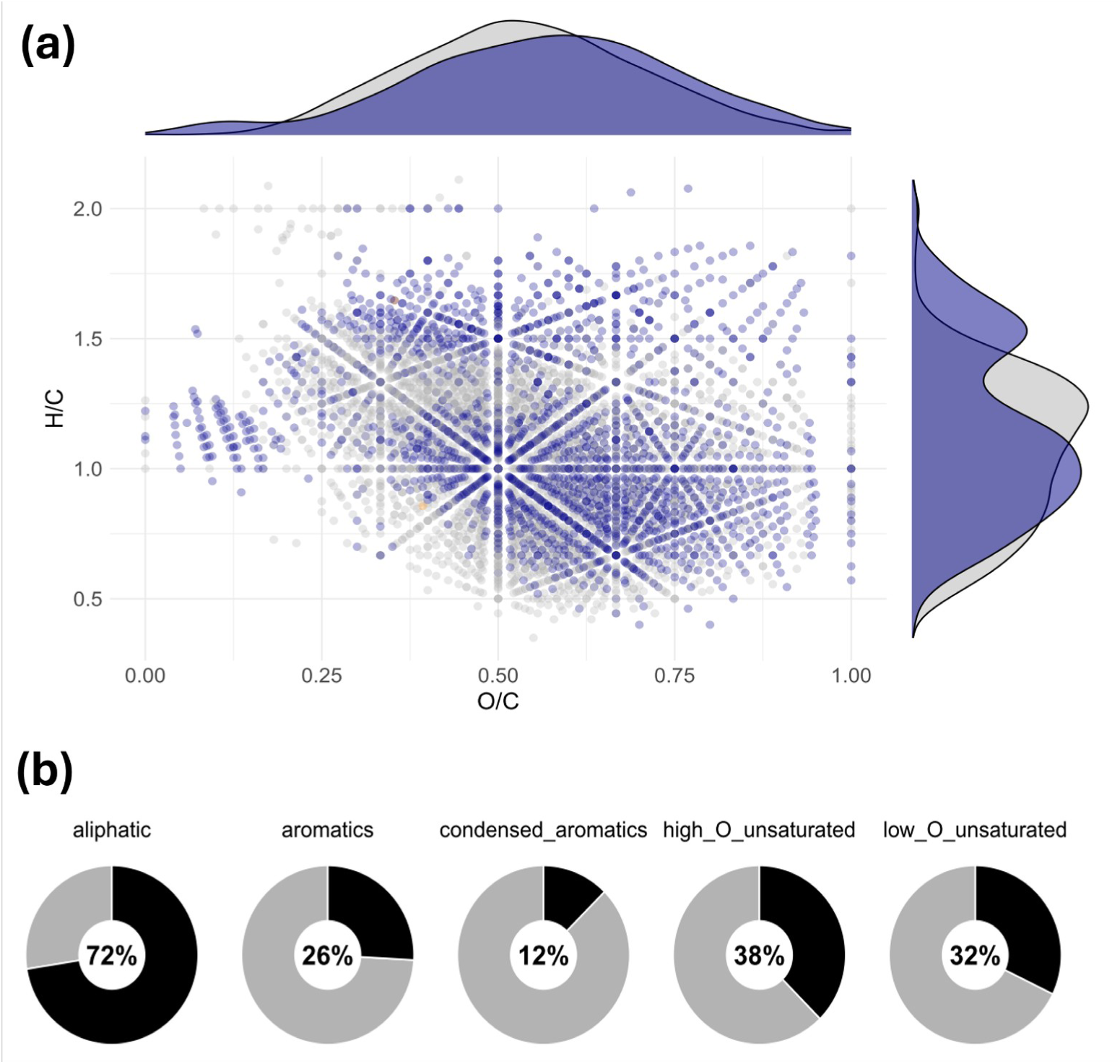
Formulas significantly correlated (adjusted p-value < 0.05) with organic carbon concentration in extracts before normalization. Panel (a) shows a Van Krevelen diagram (dark blue = positive correlation, light orange = negative correlation), panel (b) shows the percentage of correlated formulas in each chemical class. We performed Pearson correlation and adjusted p-values with the Benjamini-Hochberg method to correct for the false discovery rate.

**Figure S3.**
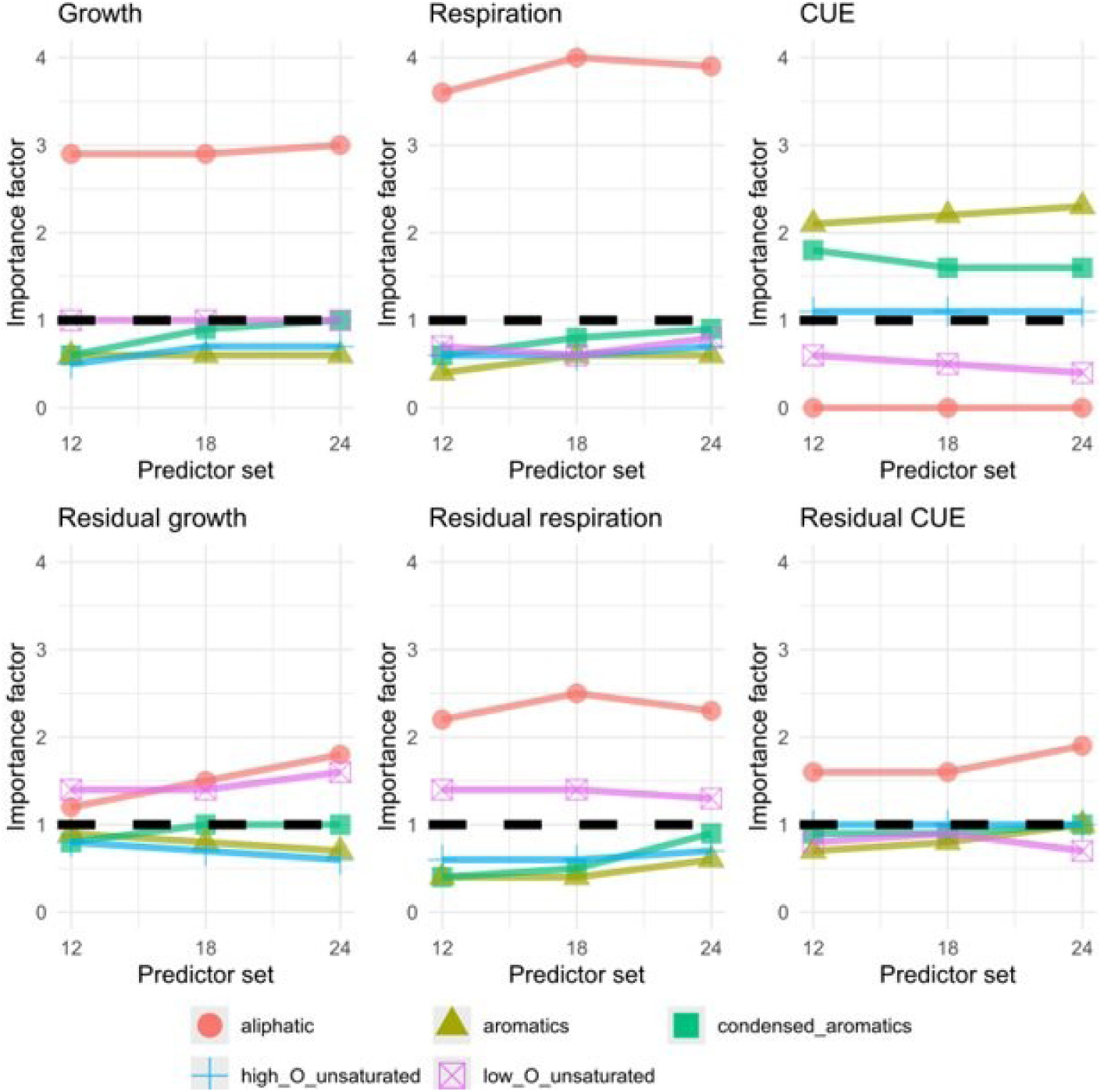
Sensitivity analysis to assess whether the study results are affected by the selection of formulas included in the PLS regression. Each panel shows the results for one response variable. The x-axis shows results for models that consider all molecular formulas that occurred in >11, >17 and >23 soils, with the 10% most important formulas retained.

**Figure S4.**
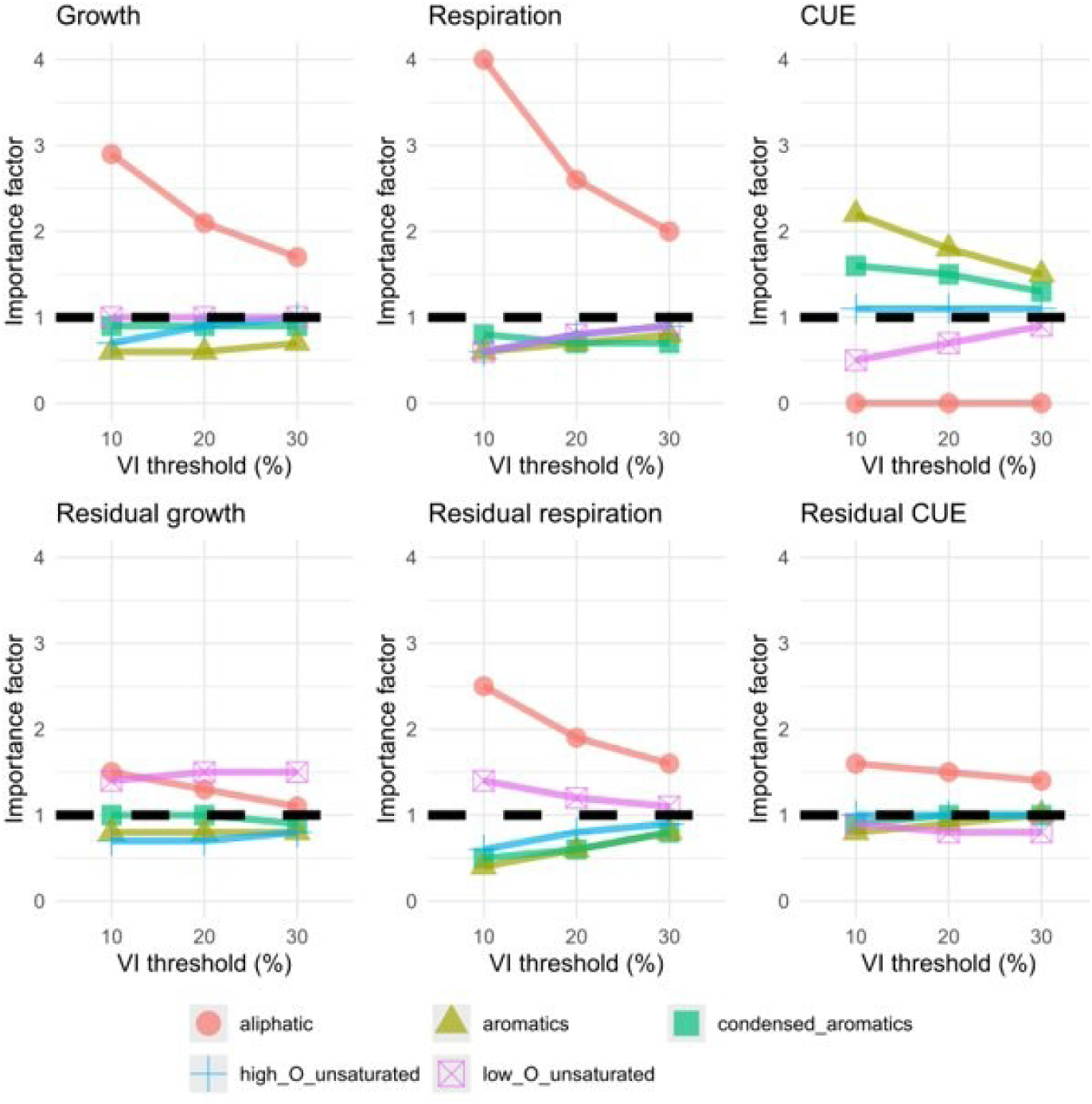
Sensitivity analysis to assess whether the study results depend on the threshold used to select predictor variables based on their variable importance (VI). Each panel shows the results for one response variable. The x-axis shows results when the 10%, 20% and 30% most important variables are interpreted, based on models that considered molecular formulas that occurred in >17 soils.

**Figure S5.**
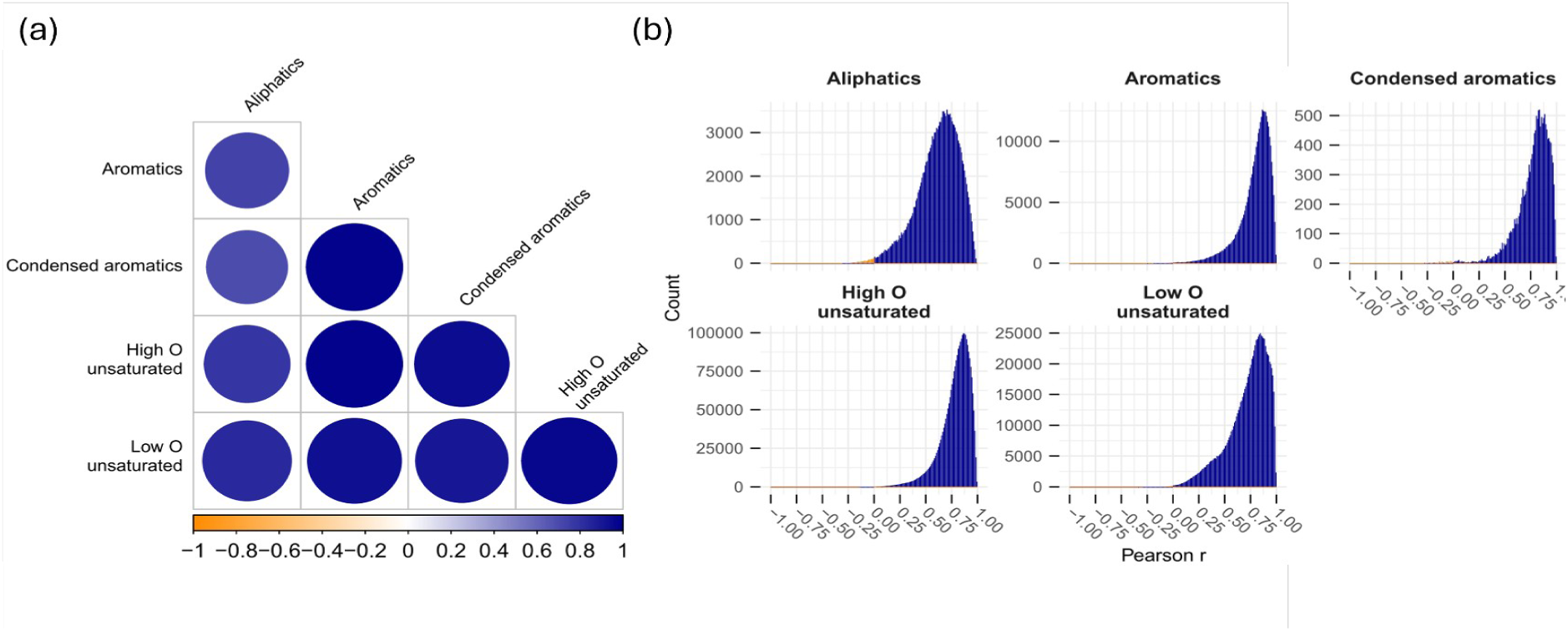
Correlation among normalized peak intensities across chemical classes for the subset of the mass spectra that was used for PLS regression. Panel (a) shows significant correlations (Pearson correlation, *p* < 0.05) between mean normalized peak intensities of the different chemical classes. Circle size and color indicate the strength of the relationships (Pearson correlation coefficient). Panel (b) shows the distribution of pairwise Pearson correlation coefficients across all compounds within each chemical class.

**Figure S6.**
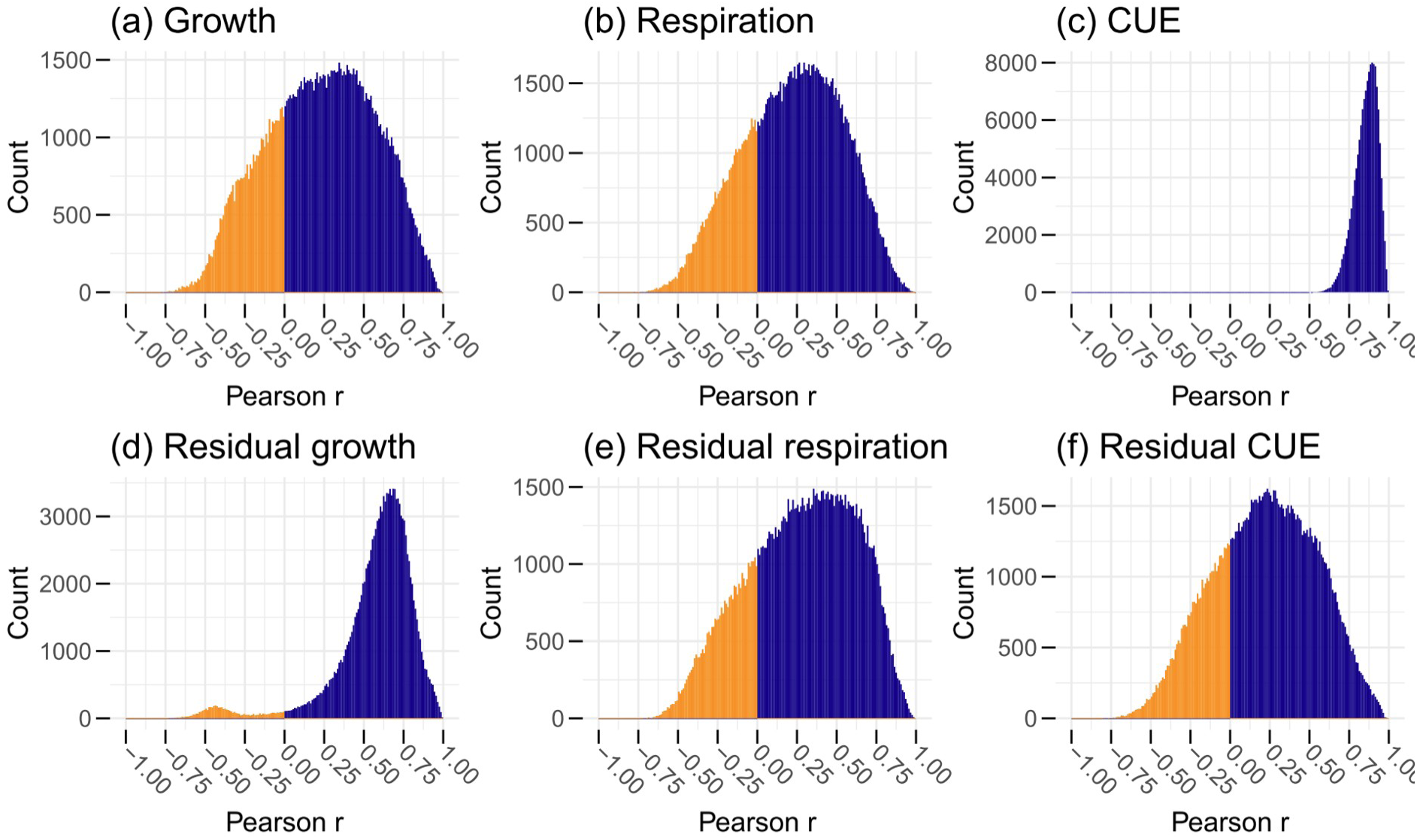
Correlation among normalized peak intensities of the top 10% most important predictive molecular formulas in each model. Shown are the distributions of pairwise Pearson correlation coefficients.

**Figure S7.**
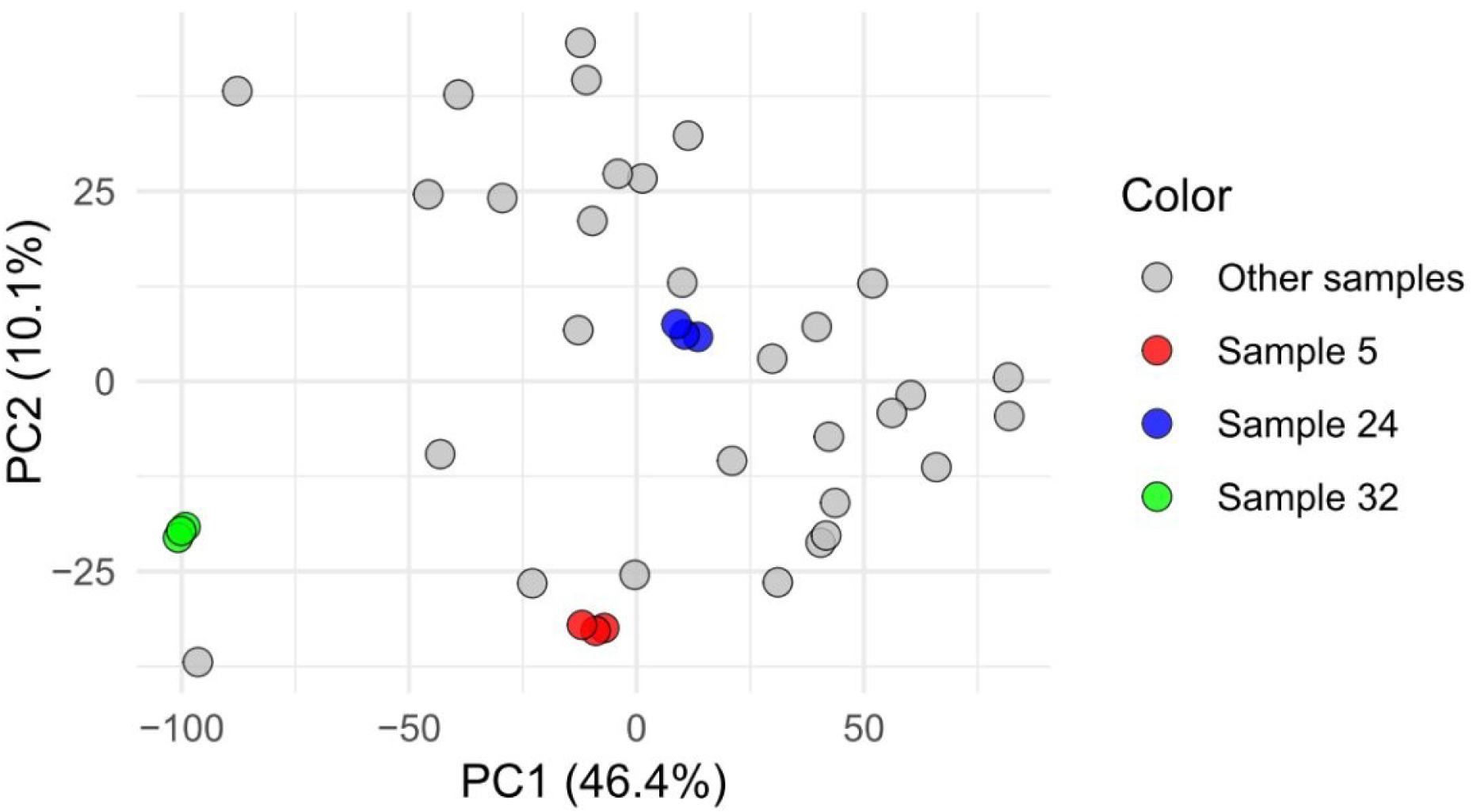
The first two principal components (together 56.5 % variation) based on the scaled signal intensities of the 5305 compounds that were retained for PLS regression. Each dot represents one sample, highlighted in colors are the three triplicates. This plot shows that the chosen methodology to characterize the chemical composition of extracted organic matter is highly reproducible and sensitive enough to reliably distinguish different samples.

**Table S1.** Summary of the chemical composition of extractable organic matter. Shown are minimum (min.), mean, median and maximum (max.) values across the 33 investigated soils, for all chemical classes together as well as for the individual chemical classes. Shown is data from the entire mass spectra (i.e., including rare molecular formulas not considered for PLS regression). °Percent of summed signal intensities per sample; *Intensity-weighted mean. NOSC = Nominal oxidation state of carbon.

| Chemical class | Parameter | Min. | Mean | Median | Max. |
| --- | --- | --- | --- | --- | --- |
| All | Number formulas | 3275 | 6855 | 6650 | 12473 |
|  | Signal intensity° (%) | 100.0 | 100.0 | 100.0 | 100.0 |
|  | Mean NOSC* | -0.22 | 0.04 | 0.05 | 0.27 |
| High O<br>Unsaturated | Number formulas | 934 | 2759 | 2783 | 5636 |
|  | Signal intensity° (%) | 29.9 | 45.6 | 46.1 | 59.7 |
|  | Mean NOSC* | 0.31 | 0.38 | 0.38 | 0.47 |
| Low O<br>Unsaturated | Number formulas | 871 | 1704 | 1742 | 3164 |
|  | Signal intensity° (%) | 18.2 | 27.2 | 26.6 | 34.1 |
|  | Mean NOSC* | -0.54 | -0.44 | -0.44 | -0.36 |
| Aliphatics | Number formulas | 635 | 1080 | 941 | 2103 |
|  | Signal intensity° (%) | 4.6 | 13.5 | 12.2 | 29.0 |
|  | Mean NOSC* | -1.02 | -0.66 | -0.65 | -0.42 |
| Aromatics | Number formulas | 354 | 986 | 943 | 1940 |
|  | Signal intensity° (%) | 8.5 | 11.3 | 10.9 | 18.4 |
|  | Mean NOSC* | 0.38 | 0.52 | 0.53 | 0.60 |
| Condensed<br>Aromatics | Number formulas | 114 | 326 | 261 | 984 |
|  | Signal intensity° (%) | 1.4 | 2.4 | 2.0 | 6.3 |
|  | Mean NOSC* | 0.67 | 0.72 | 0.71 | 0.79 |

**Table S2.** Summary of linear regression with intensity-weighted NOSC for all six response variables. All models have 31 degrees of freedom. transf. = response variable transformed with natural logarithm; coef. = coefficient.

| Variable | transf. | p-value | adj. R2 | coef. |
| --- | --- | --- | --- | --- |
| Growth | yes | 0.71 | -0.03 | 0.28 |
| Respiration | yes | 0.07 | 0.07 | -1.82 |
| **CUE** | **yes** | **0.03** | **0.12** | **1.49** |
| Residual growth | no | 0.36 | 0 | 0.45 |
| Residual respiration | no | 0.45 | -0.01 | -0.34 |
| Residual CUE | no | 0.78 | -0.03 | -0.03 |

**Table S3.** Results from two redundancy analyses to find links between organic matter composition and (a) environmental conditions and (b) microbial community composition. Predictor variables were taken from Wasner et al. (2024, Global Change Biology). Shown are significant predictors (p-value < 0.05), and percentages of total variation explained by the predictors. The “PCs” in panel (b) refer to different dimensions of microbial community composition which are explained in Wasner et al. (2024, Global Change Biology). Abbreviations: MAT = Mean annual temperature; OM = organic matter; var. = variation.

**(a) RDA with environmental condition**
| Predictor | Percent of total var. explained | Cumulative percent of total var. explained | P-value |
| --- | --- | --- | --- |
| **MAT** | 11.2 | 11.2 | 0.001 |
| **OM Quantity** | 7.9 | 19.1 | 0.014 |
| **Sand** | 3.8 | 22.9 | 0.031 |
| **pH** | 2.4 | 25.3 | 0.038 |

**(b) RDA with microbial community composition**
| Predictor | Percent of total var. explained | Cumulative percent of total var. explained | P-value |
| --- | --- | --- | --- |
| **PC1** | 17.3 | 17.3 | 0.001 |
| **PC4** | 9.5 | 26.8 | 0.002 |
| **PC2** | 6.5 | 33.3 | 0.001 |
| **PC5** | 3.0 | 36.2 | 0.014 |
| **PC8** | 2.7 | 38.9 | 0.028 |
| **PC3** | 1.4 | 40.4 | 0.035 |

## References

1. Amenabar, M.J., Shock, E.L., Roden, E.E., Peters, J.W. and Boyd, E.S., 2017. Microbial substrate preference dictated by energy demand rather than supply. Nature Geoscience, 10(8), pp.577–581. 10.1038/ngeo2978.

2. Atiwesh, G., Parrish, C.C., Banoub, J. and Le, T.T., 2022. Lignin degradation by microorganisms: A review. Biotechnol Prog., 38(2). 10.1002/btpr.3226.

3. Bajracharya, B.M., Smeaton, C.M., Markelov, I., Markelova, E., Lu, C., Cirpka, O.A. and Cappellen, P.V., 2022. Organic Matter Degradation in Energy-Limited Subsurface Environments—A Bioenergetics-Informed Modeling Approach. Geomicrobiology Journal, 39(1), pp.1–16. 10.1080/01490451.2021.1998256.

4. Bird, M.I., Wynn, J.G., Saiz, G., Wurster, C.M. and McBeath, A., 2015. The Pyrogenic Carbon Cycle. Annual Review Earth and Planetary Sciences, 43, pp.273–298.

5. Boye, K., Herrmann, A.M., Schaefer, M.V., Tfaily, M.M. and Fendorf, S., 2018. Discerning Microbially Mediated Processes During Redox Transitions in Flooded Soils Using Carbon and Energy Balances. Frontiers in Environmental Science, 6, p.15. 10.3389/fenvs.2018.00015.

6. Boye, K., Noël, V., Tfaily, M.M., Bone, S.E., Williams, K.H., Bargar, J.R. and Fendorf, S., 2017. Thermodynamically controlled preservation of organic carbon in floodplains. Nature Geoscience, 10(6), pp.415–419. 10.1038/ngeo2940.

7. Campbell, T.P., Ulrich, D.E.M., Toyoda, J., Thompson, J., Munsky, B., Albright, M.B.N., Bailey, V.L., Tfaily, M.M. and Dunbar, J., 2022. Microbial Communities Influence Soil Dissolved Organic Carbon Concentration by Altering Metabolite Composition. Frontiers in Microbiology, 12, p.799014. 10.3389/fmicb.2021.799014.

8. Carbone, M.S., Seyednasrollah, B., Rademacher, T.T., Basler, D., Le Moine, J.M., Beals, S., Beasley, J., Greene, A., Kelroy, J. and Richardson, A.D., 2019. Flux Puppy – An open-source software application and portable system design for low-cost manual measurements of CO2 and H2O fluxes. Agricultural and Forest Meteorology, 274, pp.1–6. 10.1016/j.agrformet.2019.04.012.

9. Cederlund, H., Wessén, E., Enwall, K., Jones, C.M., Juhanson, J., Pell, M., Philippot, L. and Hallin, S., 2014. Soil carbon quality and nitrogen fertilization structure bacterial communities with predictable responses of major bacterial phyla. Applied Soil Ecology, 84, pp.62–68. 10.1016/j.apsoil.2014.06.003.

10. Chakrawal, A., Calabrese, S., Herrmann, A.M. and Manzoni, S., 2022. Interacting Bioenergetic and Stoichiometric Controls on Microbial Growth. Frontiers in Microbiology, 13, p.859063. 10.3389/fmicb.2022.859063.

11. Chakrawal, A., Herrmann, A.M., Šantrůčková, H. and Manzoni, S., 2020. Quantifying microbial metabolism in soils using calorespirometry — A bioenergetics perspective. Soil Biology and Biochemistry, 148, p.107945. 10.1016/j.soilbio.2020.107945.

12. Chakrawal, A., Qafoku, O., Karra, S., Bargar, J.R. and Graham, E.B., 2025. Challenges in integrating dissolved organic matter chemodiversity into kinetic models of soil respiration. Soil Biology and Biochemistry, 211, p.109954. 10.1016/j.soilbio.2025.109954.

13. Chandel, A.K., Jiang, L. and Luo, Y., 2023. Microbial Models for Simulating Soil Carbon Dynamics: A Review. Journal of Geophysical Research: Biogeosciences, 128(8), p.e2023JG007436. 10.1029/2023JG007436.

14. Chantigny, M.H., Olk, D.C. and Angers, D.A., 2025. Investigating the nature of soil carbohydrates and amino compounds with liquid chromatography. Soil Science Society of America Journal, 89(1), p.e270018. 10.1002/saj2.70018.

15. Cyle, K.T., Klein, A.R., Aristilde, L. and Martínez, C.E., 2020. Ecophysiological Study of *Paraburkholderia* sp. Strain 1N under Soil Solution Conditions: Dynamic Substrate Preferences and Characterization of Carbon Use Efficiency. Applied and Environmental Microbiology, 86(24), pp.e01851–20. 10.1128/AEM.01851-20.

16. Deng, S., Zheng, X., Chen, X., Zheng, S., He, X., Ge, T., Kuzyakov, Y., Wu, J., Su, Y. and Hu, Y., 2021. Divergent mineralization of hydrophilic and hydrophobic organic substrates and their priming effect in soils depending on their preferential utilization by bacteria and fungi. Biology and Fertility of Soils, 57(1), pp.65–76. 10.1007/s00374-020-01503-7.

17. Doetterl, S., Stevens, A., Six, J., Merckx, R., Van Oost, K., Casanova Pinto, M., Casanova-Katny, A., Muñoz, C., Boudin, M., Zagal Venegas, E. and Boeckx, P., 2015. Soil carbon storage controlled by interactions between geochemistry and climate. Nature Geoscience, 8(10), pp.780–783. 10.1038/ngeo2516.

18. Domeignoz-Horta, L.A., Shinfuku, M., Junier, P., Poirier, S., Verrecchia, E., Sebag, D. and DeAngelis, K.M., 2021. Direct evidence for the role of microbial community composition in the formation of soil organic matter composition and persistence. ISME Communications, 1(1), p.64. 10.1038/s43705-021-00071-7.

19. Dufour, L.J.P., Herrmann, A.M., Leloup, J., Przybylski, C., Foti, L., Abbadie, L. and Nunan, N., 2022. Potential energetic return on investment positively correlated with overall soil microbial activity. Soil Biology and Biochemistry, 173, p.108800. 10.1016/j.soilbio.2022.108800.

20. Fierer, N., 2017. Embracing the unknown: disentangling the complexities of the soil microbiome. Nature Reviews Microbiology, 15(10), pp.579–590. 10.1038/nrmicro.2017.87.

21. Flamholz, A.I., Goyal, A., Fischer, W.W., Newman, D.K. and Phillips, R., 2025. The proteome is a terminal electron acceptor. Proceedings of the National Academy of Sciences, 122(1), p.e2404048121. 10.1073/pnas.2404048121.

22. Freeman, E.C., Emilson, E.J.S., Dittmar, T., Braga, L.P.P., Emilson, C.E., Goldhammer, T., Martineau, C., Singer, G. and Tanentzap, A.J., 2024. Universal microbial reworking of dissolved organic matter along environmental gradients. Nature Communications, 15(1), p.187. 10.1038/s41467-023-44431-4.

23. Friedlingstein, P., O’Sullivan, M., Jones, M.W., Andrew, R.M., Bakker, D.C.E., Hauck, J., Landschützer, P., Le Quéré, C., Luijkx, I.T., Peters, G.P., Peters, W., Pongratz, J., Schwingshackl, C., Sitch, S., Canadell, J.G., Ciais, P., Jackson, R.B., Alin, S.R., Anthoni, P., Barbero, L., Bates, N.R., Becker, M., Bellouin, N., Decharme, B., Bopp, L., Brasika, I.B.M., Cadule, P., Chamberlain, M.A., Chandra, N., Chau, T.-T.-T., Chevallier, F., Chini, L.P., Cronin, M., Dou, X., Enyo, K., Evans, W., Falk, S., Feely, R.A., Feng, L., Ford, D.J., Gasser, T., Ghattas, J., Gkritzalis, T., Grassi, G., Gregor, L., Gruber, N., Gürses, Ö., Harris, I., Hefner, M., Heinke, J., Houghton, R.A., Hurtt, G.C., Iida, Y., Ilyina, T., Jacobson, A.R., Jain, A., Jarníková, T., Jersild, A., Jiang, F., Jin, Z., Joos, F., Kato, E., Keeling, R.F., Kennedy, D., Klein Goldewijk, K., Knauer, J., Korsbakken, J.I., Körtzinger, A., Lan, X., Lefèvre, N., Li, H., Liu, J., Liu, Z., Ma, L., Marland, G., Mayot, N., McGuire, P.C., McKinley, G.A., Meyer, G., Morgan, E.J., Munro, D.R., Nakaoka, S.-I., Niwa, Y., O’Brien, K.M., Olsen, A., Omar, A.M., Ono, T., Paulsen, M., Pierrot, D., Pocock, K., Poulter, B., Powis, C.M., Rehder, G., Resplandy, L., Robertson, E., Rödenbeck, C., Rosan, T.M., Schwinger, J., Séférian, R., Smallman, T.L., Smith, S.M., Sospedra-Alfonso, R., Sun, Q., Sutton, A.J., Sweeney, C., Takao, S., Tans, P.P., Tian, H., Tilbrook, B., Tsujino, H., Tubiello, F., Van Der Werf, G.R., Van Ooijen, E., Wanninkhof, R., Watanabe, M., Wimart-Rousseau, C., Yang, D., Yang, X., Yuan, W., Yue, X., Zaehle, S., Zeng, J. and Zheng, B., 2023. Global Carbon Budget 2023. Earth System Science Data, 15(12), pp.5301–5369. 10.5194/essd-15-5301-2023.

24. Gao, D., Bai, E., Wasner, D. and Hagedorn, F., 2024. Global prediction of soil microbial growth rates and carbon use efficiency based on the metabolic theory of ecology. Soil Biology and Biochemistry, 190, p.109315. 10.1016/j.soilbio.2024.109315.

25. Garayburu-Caruso, V.A., Stegen, J.C., Song, H.-S., Renteria, L., Wells, J., Garcia, W., Resch, C.T., Goldman, A.E., Chu, R.K., Toyoda, J. and Graham, E.B., 2020a. Carbon Limitation Leads to Thermodynamic Regulation of Aerobic Metabolism. Environmental Science & Technology Letters, 7(7), pp.517–524. 10.1021/acs.estlett.0c00258.

26. Garayburu-Caruso, V.A., Stegen, J.C., Song, H.-S., Renteria, L., Wells, J., Garcia, W., Resch, C.T., Goldman, A.E., Chu, R.K., Toyoda, J. and Graham, E.B., 2020b. Carbon Limitation Leads to Thermodynamic Regulation of Aerobic Metabolism. Environmental Science & Technology Letters, 7(7), pp.517–524. 10.1021/acs.estlett.0c00258.

27. Georgiou, K., Malhotra, A., Wieder, W.R., Ennis, J.H., Hartman, M.D., Sulman, B.N., Berhe, A.A., Grandy, A.S., Kyker-Snowman, E., Lajtha, K., Moore, J.A.M., Pierson, D. and Jackson, R.B., 2021. Divergent controls of soil organic carbon between observations and process-based models. Biogeochemistry, 156(1), pp.5–17. 10.1007/s10533-021-00819-2.

28. Geyer, K.M., Dijkstra, P., Sinsabaugh, R. and Frey, S.D., 2019. Clarifying the interpretation of carbon use efficiency in soil through methods comparison. Soil Biology and Biochemistry, 128, pp.79–88. 10.1016/j.soilbio.2018.09.036.

29. Graham, E.B., Knelman, J.E., Schindlbacher, A., Siciliano, S., Breulmann, M., Yannarell, A., Beman, J.M., Abell, G., Philippot, L., Prosser, J., Foulquier, A., Yuste, J.C., Glanville, H.C., Jones, D.L., Angel, R., Salminen, J., Newton, R.J., Bürgmann, H., Ingram, L.J., Hamer, U., Siljanen, H.M.P., Peltoniemi, K., Potthast, K., Bañeras, L., Hartmann, M., Banerjee, S., Yu, R.-Q., Nogaro, G., Richter, A., Koranda, M., Castle, S.C., Goberna, M., Song, B., Chatterjee, A., Nunes, O.C., Lopes, A.R., Cao, Y., Kaisermann, A., Hallin, S., Strickland, M.S., Garcia-Pausas, J., Barba, J., Kang, H., Isobe, K., Papaspyrou, S., Pastorelli, R., Lagomarsino, A., Lindström, E.S., Basiliko, N. and Nemergut, D.R., 2016. Microbes as Engines of Ecosystem Function: When Does Community Structure Enhance Predictions of Ecosystem Processes? Frontiers in Microbiology, [online] 7. 10.3389/fmicb.2016.00214.

30. Grandy, A.S. and Neff, J.C., 2008. Molecular C dynamics downstream: The biochemical decomposition sequence and its impact on soil organic matter structure and function. Science of The Total Environment, 404(2–3), pp.297–307. 10.1016/j.scitotenv.2007.11.013.

31. Gunina, A. and Kuzyakov, Y., 2015. Sugars in soil and sweets for microorganisms: Review of origin, content, composition and fate. Soil Biology and Biochemistry, 90, pp.87–100. 10.1016/j.soilbio.2015.07.021.

32. Gunina, A. and Kuzyakov, Y., 2022. From energy to (soil organic) matter. Global Change Biology, 28(7), pp.2169–2182. 10.1111/gcb.16071.

33. Hall, S.J., Ye, C., Weintraub, S.R. and Hockaday, W.C., 2020. Molecular trade-offs in soil organic carbon composition at continental scale. Nature Geoscience, 13(10), pp.687–692. 10.1038/s41561-020-0634-x.

34. Han, L., Kaesler, J., Peng, C., Reemtsma, T. and Lechtenfeld, O.J., 2021. Online Counter Gradient LC-FT-ICR-MS Enables Detection of Highly Polar Natural Organic Matter Fractions. Analytical Chemistry, 93(3), pp.1740–1748. 10.1021/acs.analchem.0c04426.

35. Hashimoto, S., Ito, A. and Nishina, K., 2023. Divergent data-driven estimates of global soil respiration. Communications Earth & Environment, 4(1), p.460. 10.1038/s43247-023-01136-2.

36. Hawkes, J.A., D’Andrilli, J., Agar, J.N., Barrow, M.P., Berg, S.M., Catalán, N., Chen, H., Chu, R.K., Cole, R.B., Dittmar, T., Gavard, R., Gleixner, G., Hatcher, P.G., He, C., Hess, N.J., Hutchins, R.H.S., Ijaz, A., Jones, H.E., Kew, W., Khaksari, M., Palacio Lozano, D.C., Lv, J., Mazzoleni, L.R., Noriega-Ortega, B.E., Osterholz, H., Radoman, N., Remucal, C.K., Schmitt, N.D., Schum, S.K., Shi, Q., Simon, C., Singer, G., Sleighter, R.L., Stubbins, A., Thomas, M.J., Tolic, N., Zhang, S., Zito, P. and Podgorski, D.C., 2020. An international laboratory comparison of dissolved organic matter composition by high resolution mass spectrometry: Are we getting the same answer? Limnology and Oceanography: Methods, 18(6), pp.235–258. 10.1002/lom3.10364.

37. He, X., Abs, E., Allison, S.D., Tao, F., Huang, Y., Manzoni, S., Abramoff, R., Bruni, E., Bowring, S.P.K., Chakrawal, A., Ciais, P., Elsgaard, L., Friedlingstein, P., Georgiou, K., Hugelius, G., Holm, L.B., Li, W., Luo, Y., Marmasse, G., Nunan, N., Qiu, C., Sitch, S., Wang, Y.-P. and Goll, D.S., 2024. Emerging multiscale insights on microbial carbon use efficiency in the land carbon cycle. Nature Communications, 15(1), p.8010. 10.1038/s41467-024-52160-5.

38. Jones, A.R., Dalal, R.C., Gupta, V.V.S.R., Schmidt, S., Allen, D.E., Jacobsen, G.E., Bird, M., Grandy, A.S. and Sanderman, J., 2023. Molecular complexity and diversity of persistent soil organic matter. Soil Biology and Biochemistry, 184, p.109061. 10.1016/j.soilbio.2023.109061.

39. Jones, D.L., Hill, P.W., Smith, A.R., Farrell, M., Ge, T., Banning, N.C. and Murphy, D.V., 2018. Role of substrate supply on microbial carbon use efficiency and its role in interpreting soil microbial community-level physiological profiles (CLPP). Soil Biology and Biochemistry, 123, pp.1–6. 10.1016/j.soilbio.2018.04.014.

40. Kallenbach, C.M., Frey, S.D. and Grandy, A.S., 2016. Direct evidence for microbial-derived soil organic matter formation and its ecophysiological controls. Nature Communications, 7(1), p.13630. 10.1038/ncomms13630.

41. Kästner, M., Maskow, T., Miltner, A., Lorenz, M. and Thiele-Bruhn, S., 2024. Assessing energy fluxes and carbon use in soil as controlled by microbial activity - A thermodynamic perspective A perspective paper. Soil Biology and Biochemistry, 193, p.109403. 10.1016/j.soilbio.2024.109403.

42. Kästner, M., Miltner, A., Thiele-Bruhn, S. and Liang, C., 2021. Microbial Necromass in Soils—Linking Microbes to Soil Processes and Carbon Turnover. Frontiers in Environmental Science, 9, p.756378. 10.3389/fenvs.2021.756378.

43. Keiluweit, M., Nico, P.S., Kleber, M. and Fendorf, S., 2016. Are oxygen limitations under recognized regulators of organic carbon turnover in upland soils? Biogeochemistry, 127(2–3), pp.157–171. 10.1007/s10533-015-0180-6.

44. Khatoon, H., Solanki, P., Narayan, M., Tewari, L. and Rai, J., 2017. Role of microbes in organic carbon decomposition and maintenance of soil ecosystem. International Journal of Chemical Studies.

45. Knicker, H., 2011. Pyrogenic organic matter in soil: Its origin and occurrence, its chemistry and survival in soil environments. Quaternary International, 243(2), pp.251–263. 10.1016/j.quaint.2011.02.037.

46. Koch, B.P. and Dittmar, T., 2016. From mass to structure: an aromaticity index for high-resolution mass data of natural organic matter. Rapid Communications in Mass Spectrometry, 30(1), pp.250–250. 10.1002/rcm.7433.

47. Kögel-Knabner, I. and Amelung, W., 2014. Dynamics, Chemistry, and Preservation of Organic Matter in Soils. In: Treatise on Geochemistry. [online] Elsevier. pp.157–215. 10.1016/B978-0-08-095975-7.01012-3.

48. Kopittke, P.M., Dalal, R.C., McKenna, B.A., Smith, P., Wang, P., Weng, Z., Van Der Bom, F.J.T. and Menzies, N.W., 2024. Soil is a major contributor to global greenhouse gas emissions and climate change. SOIL, 10(2), pp.873–885. 10.5194/soil-10-873-2024.

49. Kraus, T.E.C., Dahlgren, R.A. and Zasoski, R.J., 2003. Tannins in nutrient dynamics of forest ecosystems - a review. Plant and Soil, 256(1), pp.41–66. 10.1023/A:1026206511084.

50. Kuhn, M., 2008. Building Predictive Models in R Using the caret Package. Journal of Statistical Software, 28(5), pp.1–26. 10.18637/jss.v028.i05,%2520 https://www.jstatsoft.org/index.php/jss/article/view/v028i05.

51. LaRowe, D.E. and Van Cappellen, P., 2011. Degradation of natural organic matter: A thermodynamic analysis. Geochimica et Cosmochimica Acta, 75(8), pp.2030–2042. 10.1016/j.gca.2011.01.020.

52. Laszakovits, J.R. and MacKay, A.A., 2022. Data-Based Chemical Class Regions for Van Krevelen Diagrams. Journal of the American Society for Mass Spectrometry, 33(1), pp.198– 202. 10.1021/jasms.1c00230.

53. Lechtenfeld, O.J., Kaesler, J., Jennings, E.K. and Koch, B.P., 2024. Direct Analysis of Marine Dissolved Organic Matter Using LC-FT-ICR MS. Environmental Science & Technology, 58(10), pp.4637–4647. 10.1021/acs.est.3c07219.

54. Lehmann, J., Hansel, C.M., Kaiser, C., Kleber, M., Maher, K., Manzoni, S., Nunan, N., Reichstein, M., Schimel, J.P., Torn, M.S., Wieder, W.R. and Kögel-Knabner, I., 2020. Persistence of soil organic carbon caused by functional complexity. Nature Geoscience, 13(8), pp.529–534. 10.1038/s41561-020-0612-3.

55. Lehmann, J. and Kleber, M., 2015. The contentious nature of soil organic matter. Nature, 528(7580), pp.60–68. 10.1038/nature16069.

56. Leide, J., Nierop, K.G.J., Deininger, A.-C., Staiger, S., Riederer, M. and De Leeuw, J.W., 2020. Leaf cuticle analyses: implications for the existence of cutan/non-ester cutin and its biosynthetic origin. Annals of Botany, 126(1), pp.141–162. 10.1093/aob/mcaa056.

57. Liang, C., Schimel, J.P. and Jastrow, J.D., 2017. The importance of anabolism in microbial control over soil carbon storage. Nature Microbiology, 2(8), p.17105. 10.1038/nmicrobiol.2017.105.

58. Liland, K., Mevik, B. and Wehrens, R., 2024. pls: Partial Least Squares and Principal Component Regression. 10.32614/CRAN.package.pls.

59. Lützow, M.V., Kögel-Knabner, I., Ekschmitt, K., Matzner, E., Guggenberger, G., Marschner, B. and Flessa, H., 2006. Stabilization of organic matter in temperate soils: mechanisms and their relevance under different soil conditions – a review. European Journal of Soil Science, 57(4), pp.426–445. 10.1111/j.1365-2389.2006.00809.x.

60. Mainka, M., Summerauer, L., Wasner, D., Garland, G., Griepentrog, M., Berhe, A.A. and Doetterl, S., 2022. Soil geochemistry as a driver of soil organic matter composition: insights from a soil chronosequence. Biogeosciences, 19(6), pp.1675–1689. 10.5194/bg-19-1675-2022.

61. Malik, A.A., Martiny, J.B.H., Brodie, E.L., Martiny, A.C., Treseder, K.K. and Allison, S.D., 2020. Defining trait-based microbial strategies with consequences for soil carbon cycling under climate change. The ISME Journal, 14(1), pp.1–9. 10.1038/s41396-019-0510-0.

62. Mooshammer, M., Wanek, W., Zechmeister-Boltenstern, S. and Richter, A., 2014. Stoichiometric imbalances between terrestrial decomposer communities and their resources: mechanisms and implications of microbial adaptations to their resources. *Frontiers in Microbiology*, [online] 5. 10.3389/fmicb.2014.00022.

63. Oksanen, J., Simpson, G., Blanchet, F., Kindt, R., Legendre, P., Minchin, P., ÓHara, R., Solymos, P. and, et al., 2025. vegan: Community Ecology Package. 10.32614/CRAN.package.vegan.

64. Piton, G., Allison, S.D., Bahram, M., Hildebrand, F., Martiny, J.B.H., Treseder, K.K. and Martiny, A.C., 2023. Life history strategies of soil bacterial communities across global terrestrial biomes. Nature Microbiology, 8(11), pp.2093–2102. 10.1038/s41564-023-01465-0.

65. Quiquampoix, H. and Burns, R.G., 2007. Interactions between Proteins and Soil Mineral Surfaces: Environmental and Health Consequences. Elements, 3(6), pp.401–406. 10.2113/GSELEMENTS.3.6.401.

66. Roth, V.-N., Dittmar, T., Gaupp, R. and Gleixner, G., 2015. The Molecular Composition of Dissolved Organic Matter in Forest Soils as a Function of pH and Temperature. PLOS ONE, 10(3), p.e0119188. 10.1371/journal.pone.0119188.

67. Shi, C., Mudunuru, M., Bowman, M., Zhao, Q., Toyoda, J., Kew, W., Corilo, Y., Qafoku, O., Bargar, J.R., Karra, S. and Graham, E.B., 2025. Scaling High-Resolution Soil Organic Matter Composition to Improve Predictions of Potential Soil Respiration Across the Continental United States. Geophysical Research Letters, 52(4), p.e2024GL113091. 10.1029/2024GL113091.

68. Simon, C., Miltner, A., Mulder, I., Kaiser, K., Lorenz, M., Thiele-Bruhn, S. and Lechtenfeld, O., 2025. Long-term effects of manure addition on soil organic matter molecular composition: Carbon transformation as a major driver of energetic potential. Soil Biology and Biochemistry, 205, p.109755. 10.1016/j.soilbio.2025.109755.

69. Simpson, M.J. and Simpson, A.J., 2012. The Chemical Ecology of Soil Organic Matter Molecular Constituents. Journal of Chemical Ecology, 38(6), pp.768–784. 10.1007/s10886-012-0122-x.

70. Sinsabaugh, R.L., 2010. Phenol oxidase, peroxidase and organic matter dynamics of soil. Soil Biology and Biochemistry, 42(3), pp.391–404. 10.1016/j.soilbio.2009.10.014.

71. Sokol, N.W., Slessarev, E., Marschmann, G.L., Nicolas, A., Blazewicz, S.J., Brodie, E.L., Firestone, M.K., Foley, M.M., Hestrin, R., Hungate, B.A., Koch, B.J., Stone, B.W., Sullivan, M.B., Zablocki, O., LLNL Soil Microbiome Consortium, Trubl, G., McFarlane, K., Stuart, R., Nuccio, E., Weber, P., Jiao, Y., Zavarin, M., Kimbrel, J., Morrison, K., Adhikari, D., Bhattacharaya, A., Nico, P., Tang, J., Didonato, N., Paša-Tolić, L., Greenlon, A., Sieradzki, E.T., Dijkstra, P., Schwartz, E., Sachdeva, R., Banfield, J. and Pett-Ridge, J., 2022. Life and death in the soil microbiome: how ecological processes influence biogeochemistry. Nature Reviews Microbiology, 20(7), pp.415–430. 10.1038/s41579-022-00695-z.

72. Spohn, M., Klaus, K., Wanek, W. and Richter, A., 2016. Microbial carbon use efficiency and biomass turnover times depending on soil depth – Implications for carbon cycling. Soil Biology and Biochemistry, 96, pp.74–81. 10.1016/j.soilbio.2016.01.016.

73. Stumpf, K., Simon, C., Miltner, A., Maskow, T. and Lechtenfeld, O.J., 2025. Deciphering the energy use channels in soil organic Matter: Impacts of long-term manure addition and necromass revealed by LC-FT-ICR-MS. Soil Biology and Biochemistry, 208, p.109857. 10.1016/j.soilbio.2025.109857.

74. Sulman, B.N., Moore, J.A.M., Abramoff, R., Averill, C., Kivlin, S., Georgiou, K., Sridhar, B., Hartman, M.D., Wang, G., Wieder, W.R., Bradford, M.A., Luo, Y., Mayes, M.A., Morrison, E., Riley, W.J., Salazar, A., Schimel, J.P., Tang, J. and Classen, A.T., 2018. Multiple models and experiments underscore large uncertainty in soil carbon dynamics. Biogeochemistry, 141(2), pp.109–123. 10.1007/s10533-018-0509-z.

75. Torgo, L., 2016. Data Mining with R, learning with case studies, 2nd edition. [online] Chapman and Hall/CRC. Available at: <http://ltorgo.github.io/DMwR2>.

76. Trivedi, P., Anderson, I.C. and Singh, B.K., 2013. Microbial modulators of soil carbon storage: integrating genomic and metabolic knowledge for global prediction. Trends in Microbiology, 21(12), pp.641–651. 10.1016/j.tim.2013.09.005.

77. Turner, J.W., Hartman, B.E. and Hatcher, P.G., 2013. Structural characterization of suberan isolated from river birch (Betula nigra) bark. Organic Geochemistry, 57, pp.41–53. 10.1016/j.orggeochem.2013.01.004.

78. Vance, E.D., Brookes, P.C. and Jenkinson, D.S., 1987. An extraction method for measuring soil microbial biomass C. Soil Biology and Biochemistry, 19(6), pp.703–707. 10.1016/0038-0717(87)90052-6.

79. Von Stockar, U., 2010. Biothermodynamics of live cells: a tool for biotechnology and biochemical engineering. Journal of Non-Equilibrium Thermodynamics, [online] 35(4). 10.1515/jnetdy.2010.024.

80. Wang, C. and Kuzyakov, Y., 2023. Energy use efficiency of soil microorganisms: Driven by carbon recycling and reduction. Global Change Biology, 29(22), pp.6170–6187. 10.1111/gcb.16925.

81. Wasner, D., Abramoff, R., Griepentrog, M., Venegas, E.Z., Boeckx, P. and Doetterl, S., 2024a. The Role of Climate, Mineralogy and Stable Aggregates for Soil Organic Carbon Dynamics Along a Geoclimatic Gradient. Global Biogeochemical Cycles, 38(7), p.e2023GB007934. 10.1029/2023GB007934.

82. Wasner, D., Han, X., Schnecker, J., Frossard, A., Venegas, E.Z. and Doetterl, S., 2025. Quantity Versus Quality: Links Between Soil Organic Matter and Bacterial Community Composition Along a Geoclimatic Gradient. Environmental Microbiology, 27(3), p.e70070. 10.1111/1462-2920.70070.

83. Wasner, D., Prommer, J., Zezula, D., Mooshammer, M., Hu, Y. and Wanek, W., 2023. Tracing 33P-labelled organic phosphorus compounds in two soils: New insights into decomposition dynamics and direct use by microbes. Frontiers in Soil Science, 3, p.1097965. 10.3389/fsoil.2023.1097965.

84. Wasner, D., Schnecker, J., Han, X., Sun, Y., Frossard, A., Zagal Venegas, E., Boeckx, P. and Doetterl, S., 2024b. Environment and microbiome drive different microbial traits and functions in the macroscale soil organic carbon cycle. Global Change Biology, 30(8), p.e17465. 10.1111/gcb.17465.

85. Wieder, W.R., Allison, S.D., Davidson, E.A., Georgiou, K., Hararuk, O., He, Y., Hopkins, F., Luo, Y., Smith, M.J., Sulman, B., Todd-Brown, K., Wang, Y., Xia, J. and Xu, X., 2015. Explicitly representing soil microbial processes in Earth system models. Global Biogeochemical Cycles, 29(10), pp.1782–1800. 10.1002/2015GB005188.

86. Wilcke, W., 2000. SYNOPSIS Polycyclic Aromatic Hydrocarbons (PAHs) in Soil — a Review. Journal of Plant Nutrition and Soil Science, 163(3), pp.229–248. 10.1002/1522-2624(200006)163:3<229::AID-JPLN229>3.0.CO;2-6.

87. Wurz, J., Groß, A., Franze, K. and Lechtenfeld, O., 2024. Lambda-Miner: Enhancing Reproducible Natural Organic Matter Data Processing with a Semi-Automatic Web Application. [online] EGU General Assembly 2024 . 10.5194/egusphere-egu24-15782.

88. Zeileis, A. and Hothorn, T., 2002. Diagnostic Checking in Regression Relationships. R News, 2(3), pp.7–10.

